# Mitochondrial Dual-coding Genes in *Trypanosoma* brucei

**DOI:** 10.1101/187732

**Authors:** Laura E. Kirby, Donna Koslowsky

**Affiliations:** Department of Microbiology and Molecular Genetics, Michigan State University, East Lansing, Michigan, United States of America

## Abstract

*Trypanosoma brucei* is transmitted between mammalian hosts by the tsetse fly. In the mammal, they are exclusively extracellular, continuously replicating within the bloodstream. During this stage, the mitochondrion lacks a functional electron transport chain (ETC). Successful transition to the fly, requires activation of the ETC and ATP synthesis via oxidative phosphorylation. This life cycle leads to a major problem: in the bloodstream, the mitochondrial genes are not under selection and are subject to genetic drift that endangers their integrity. Exacerbating this, *T. brucei* undergoes repeated population bottlenecks as they evade the host immune system that would create additional forces of genetic drift. These parasites possess several unique genetic features, including RNA editing of mitochondrial transcripts. RNA editing creates open reading frames by the guided insertion and deletion of U-residues within the mRNA. A major question in the field has been why this metabolically expensive system of RNA editing would evolve and persist. Here, we show that many of the edited mRNAs can alter the choice of start codon and the open reading frame by alternative editing of the 5’ end. Analyses of mutational bias indicate that six of the mitochondrial genes may be dual-coding and that RNA editing allows access to both reading frames. We hypothesize that dual-coding genes can protect genetic information by essentially hiding a non-selected gene within one that remains under selection. Thus, the complex RNA editing system found in the mitochondria of trypanosomes provides a unique molecular strategy to combat genetic drift in non-selective conditions.

**Author Summary:** In African trypanosomes, many of the mitochondrial mRNAs require extensive RNA editing before they can be translated. During this process, each edited transcript can undergo hundreds of cleavage/ligation events as U-residues are inserted or deleted to generate a translatable open reading frame. A major paradox has been why this incredibly metabolically expensive process would evolve and persist. In this work, we show that many of the mitochondrial genes in trypanosomes are dual-coding, utilizing different reading frames to potentially produce two very different proteins. Access to both reading frames is made possible by alternative editing of the 5’ end of the transcript. We hypothesize that dual-coding genes may work to protect the mitochondrial genes from mutations during growth in the mammalian host, when many of the mitochondrial genes are not being used. Thus, the complex RNA editing system may be maintained because it provides a unique molecular strategy to combat genetic drift.

## Introduction

Trypanosomes are one of the most successful parasites in existence, inhabiting an incredibly wide range of hosts [1, 2]. The dixenous members cycle between two distinct hosts and can encounter different environments with distinct metabolic constraints. These parasites are unique in that they all possess glycosomes (where glycolysis occurs) as well as mitochondria [3]. The salivarian trypanosomes (e.g. *T. brucei*, *T. vivax*) are especially interesting, because they are exclusively extracellular in their mammalian hosts, continuously replicating within the bloodstream over periods of months. During this stage of the life cycle, the mitochondrion is down-regulated, lacking both Krebs cycle enzymes and a functional electron transport chain (ETC) [4]. Successful transition to the fly vector, requires activation of the ETC and ATP synthesis via oxidative phosphorylation. This unique lifecycle leads to a major problem: when the mitochondrial genes are unused, they are not under selection, hence the integrity of these genes are threatened by genetic drift [5, 6]. Exacerbating this, salivarian trypanosomes undergo a severe bottleneck as they transition through the tsetse fly and into the mammalian host, and then within the bloodstream, they undergo multiple bottlenecks at each antigenic switch, as they evade the host immune system [7]. Such bottlenecks create additional forces of genetic drift, where genes can be lost even if their deleterious fitness effect is considerable. These parasites possess several unique genetic features, including RNA editing of the mitochondrial transcripts. RNA editing creates open reading frames in “cryptogenes” by insertion and deletion of uridylate residues at specific sites within the mRNA. The U-insertions/deletions are directed by small guide RNAs (gRNA) and can repair frameshifts, generate start and stop codons and more than double the size of the transcript (for review see [8]). While the mRNA cryptogenes are encoded on maxicircles (25-50 copies per DNA network), the guide RNAs are encoded on thousands of 1 kb minicircles, encoding 3 – 5 gRNA genes each [9]. This effectively means that the genetic information for the edited mitochondrial mRNAs is dispersed between the mRNA cryptogenes on the maxicircles and the thousands of gRNA coding minicircles. The extensive editing of a single transcript can require more than 40 gRNAs and hundreds of editing events [10]. While the initial gRNA can interact with the 3’ end of the pre-edited transcript, all subsequent gRNAs anchor to edited sequence created by the preceding gRNA. Hence, editing proceeds from the 3’ end to the 5’ end of the mRNA transcript with the terminal gRNA (last one in the cascade) often creating the start codon needed for translation. This sequential dependence means that with even high accuracy rates for each gRNA, the overall fidelity of the process is astonishingly low. A major question in the field has been why this fragile and metabolically expensive system of RNA editing would evolve and persist.

Another level of complexity in the kinetoplastids RNA editing process was the detection of an alternative editing event that leads to the production of a functionally discrete protein isoform. Alternative editing of Cytochrome Oxidase III (*COIII*) is reported to generate a novel DNA-binding protein, AEP-1, that functions in mitochondrial DNA maintenance [11, 12]. In this transcript, one alternative gRNA generates sequence changes at two sites that links an open reading frame (ORF) found in the pre-edited 5’ end, to the 3’ transmembrane domains found in the *COIII* edited ORF. This was the first indication, that one cryptogene could contain information for more than one protein. Here, we show that as many as six additional cryptogenes also encode for more than one protein. Analyses of the terminal gRNA populations indicate that gRNA sequence variants exist that can alter the choice of the start codon and the open reading frame by alternative editing of the 5’ end of the mRNA. Mutational bias analyses indicate that six of the mitochondrial genes may be dual-coding, with RNA editing allowing access to both reading frames. Dual-coding genes are defined as a stretch of DNA containing overlapping open reading frames (ORFs) [13, 14]. Of particular interest are dual-coding genes that contain two ORFs read in the same direction: a canonical protein (normally annotated as protein coding in the literature) and an alternative ORF. Maintaining dual-coding genes is costly, as it constrains the flexibility of the amino acid composition of both proteins. Hence, it is thought that dual-coding genes can survive long evolutionary spans only if the overlap is advantageous to the organism [15]. We hypothesize that trypanosomes use dual-coding genes to protect genetic information by essentially hiding a non-selected (ETC) gene within one that remains under selection. Thus, the ability to access overlapping reading frames may be added to a growing list of gene protective strategies made possible by the complex RNA editing process [5, 6, 16].

## Materials and Methods

### Trypanosome growth

*T. brucei* procyclic clones from IsTAR (Eatro 164), TREU 667 and TREU 927 cell lines were grown in SDM79 at 27 °C and harvested at a cell density of 1-3x10^7^. The TREU 667 cell line was originally isolated from a bovine host in 1966 in Uganda [17]. The TREU 927 cell line was originally isolated from *Glossina pallidipes* in 1970 in Kenya [18]. The Eatro 164 strain was isolated in 1960 from *Alcephalus lictensteini* and maintained in the lab of Dr. K. Vickerman until being obtained by Dr. Ken Stuart in 1966 [19]. Dr. Stuart derived the procyclic form from the bloodstream form culture in 1979.

### Guide RNA isolation, preparation, and Sequencing

Mitochondrial mRNAs and gRNAs were isolated as previously described [10]. All RNAs were treated with Promega DNAse RQI. In order to isolate gRNAs from TREU 667 and TREU 927 cells, RNAs were size fractionated on a polyacrylamide gel as previously described [10]. gRNAs were then extracted and prepped for sequencing using the Illumina Small RNA protocol [10]. Libraries from TREU 667 and TREU 927 were deep sequenced on the Illumina GAIIx; reads were processed and trimmed as previously described [10].

### Messenger RNA Preparation and Sequencing

In order to isolate target mRNAs, isolated TREU 667 mitochondrial RNAs were reverse transcribed using the Applied Biosystems High Capacity cDNA Reverse Transcription Kit. *CR3* cDNAs were amplified via PCR using the following primers (underlined portions are gene specific and non-underlined portions are tag regions used in deep sequencing reaction):

> CR3DS5´NEV: ACACTGACGACATGGTTCTAC<u>AAGAAATATAAATATGTGTATG</u>
>
> CR3DS3´170: TACGGTAGCAGAGACTTGGTCT<u>CAATAAACCCATATTAAATAAAAAACAAAAATCC</u>

After amplification, the products were purified using the QIAquick PCR Purification Kit, and paired end Illumina deep sequencing was performed on the Illumina Miseq (2x 250 bp paired end run). Low quality results were removed using FaQCs, adapters were removed using Trimmomatic and PEAR was used to merge paired end reads. Finally, Fastx was used to compile identical reads while maintaining the number of redundant reads. *CR3* edited transcripts were identified by comparing sequence downstream of the 5’ never edited region to the edited *CR3* sequence. gRNAs were identified by using the mRNA sequences as queries against our existing gRNA databases, as previously described [10].

### Mutational Frequency and Editing Conservation Analysis

Mitochondrial pan-edited genes were categorized as potentially dual-coding based on identification of extended alternative reading frames and/or presence of identified gRNAs that generate alternative 5’ end sequences. These genes include *CR3*, *CR4*, *ND3*, the 5’ editing domain of *ND7*, *ND9* and *RPS12*. Nondual-coding pan-edited genes include ATPase 6, COIII, ND8 and the 3’ editing domain of *ND7*. Partially edited genes include *CYb*, *Murf II* and *COII*. Never edited genes include *COI*, *ND1*, *ND2*, *ND4* and *ND5*. For all analyses, *ND7* was considered as two separate coding regions: the 5’ editing domain (*ND7N*) and the 3’ editing domain (*ND7C*) [20]. As we hypothesize that only the 5’ editing domain of *ND7* is dual-coding, mutation calculations for *ND7N* was pooled with the dual-coding genes and *ND7C* was pooled with nondual-coding pan-edited genes. *T. brucei* and *T. vivax* mRNA sequences of mitochondrial encoded genes were aligned based on protein sequence using Clustal Omega [21]. Nucleotide sequence mutations were identified and their effects on the amino acid sequence were classified as silent, missense or nonsense mutations. Missense mutations were further divided into three groups based on the PAM 250 matrix where conversions with a value <0 were considered not conserved, conversions with a value 0≤x≤0.5 were considered modestly conserved, and conversions with a value >0.5 were considered strongly conserved [22]. Mutation frequencies were normalized for each gene using nucleotide sequence length. Frequencies were compared using unpaired t-tests.

The extent of editing conservation between *T. vivax* and *T. brucei* was calculated by aligning the pan-edited genes based on ACG sequence. For each alignment, each location between an A, C or G nucleotide where a U-residue was inserted or deleted in either sequence was considered an editing site. Editing sites were classified as identical in both sequences, altered in insertion or deletion length, having switched from an insertion site to a deletion site, or only occurring in one of the sequences. Percent editing conservation was based on total number of editing sites within each mRNA. Percentages were compared using unpaired t-tests.

A principal component analysis (PCA) was performed on all three reading frames of the pan-edited genes using the scikit-learn principal component analysis tool [23]. For this analysis, the predicted protein sequences for all three reading frames were aligned using Clustal Omega [21]. Missense, nonsense, and indel mutations were quantified. Missense mutations were further divided into three groups as described above. Each mutation type was quantified and the relative frequency of each mutation calculated based on protein amino acid length. The variables used in the PCA include the protein mutation frequencies and the percentage of identical editing sites in each mRNA. The first reading frame of each gene is defined as the ORF published in the literature.

### Data and Software Availability

CR3 sequence accession number: SAMN06318039. TREU 927 gRNA sequence accession number: SAMN06318154. TREU 667 gRNA sequence accession number: SAMN06318153. NCBI’s Sequence Read Archive. Code used in PCA available upon request.

## Results

In *T. brucei*, analyses of the gRNA transcriptome for the pan-edited transcripts indicate that full editing involves a large number of gRNA populations [10, 24]. In addition, most of the gRNA populations (population defined as guiding the same or near same region of the mRNA) contain multiple sequence classes. The sequence classes most often differ in R to R or Y to Y mutations, hence guide the generation of the same mRNA sequence (A:U and G:U basepairs allowed). During these analyses, we noted that the terminal gRNA population for Cytosine-rich Region 3 (*CR3*) (putative NADH dehydrogenase subunit 4L [25]), had 3’ sequences that would extend editing beyond the previously identified translation start codon. In addition, this population had several sequence variants that would generate different edited sequences in this region. The most abundant terminal gRNA would introduce a stop codon in-frame with two alternative AUG start codons found near the 5’ end (Fig 1A). Other sequence classes however, would either bring the upstream AUGs into frame, or shift the reading frame. Intriguingly, the alternative +1 reading frame (ARF) did not contain any premature termination codons. In order to determine if these gRNAs were utilized, we used Illumina deep sequencing to identify the most abundant forms of fully edited *CR3* transcripts. Surprisingly, we identified multiple forms of the mRNA (Fig 1 A,B,C and S1 Table). The first was the fully edited sequence predicted by the most abundant gRNA identified (Fig 1 A). The other transcripts however, had unique editing patterns at the 5’ end (Fig 1B,C and S1 Table). Use of these 5’ *CR3* sequences allowed us to identify novel gRNAs. Predicted translation of these mRNA sequences indicate that they use the +1 reading frame, and that the protein generated would be the same length as the ORF previously identified. This suggests that *CR3* is dual-coding, and that selection of the terminal gRNA determines which reading frame will be used. A re-examination of the terminal gRNAs for the pan-edited genes indicated that at least two other transcripts, NADH dehydrogenase subunit 7 (*ND7*) and ribosomal protein subunit 12 (*RPS12*), have identified gRNA sequence variants within the terminal gRNA population that allow access to alternative reading frames (Fig 1 D – E and S2 and S3 Tables). Interestingly, the alternative gRNA for *ND7* generates a +2 frameshift with a 65 amino acid open reading frame. The *ND7* transcript is differentially edited in two distinct domains separated by 59 nts that are not edited in the mature transcript (the HR3 region) [20]. Only the 5’ domain is edited in both life cycle stages, full editing of the 3’ domain was only found in bloodstream form (BF) parasites. The stop codon for the +2 frameshift is found within the HR3 region, therefore this alternative protein would be generated by full editing of only the 5’ domain. While the most abundant gRNA in the Eatro BF transcriptome (~50,000 reads) would generate a sequence utilizing the identified *ND7* ORF (Fig 1 C ORF), the most abundant gRNA ( >100,000) in the Eatro 164 procyclic library is the +2 ARF gRNA (Fig 1 C ARF and S2 Table).

**Fig 1.**
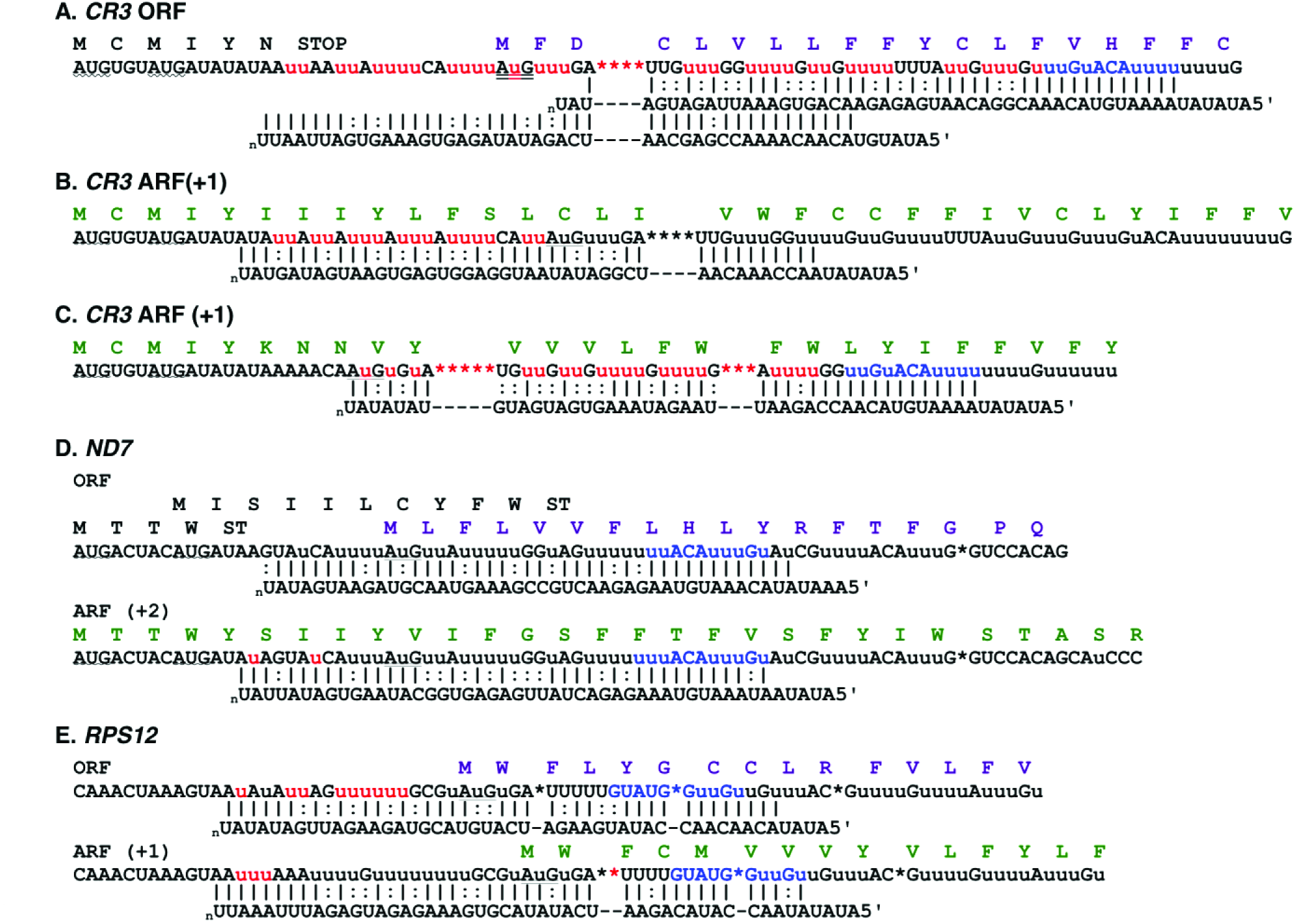
Alternative editing of the 5’ end of pan-edited genes results in access to different reading frames. *CR3* (A,B,C): Sequenced mRNA variants are aligned with gRNAs and predicted protein sequences. Inserted U-residues are lowercase while deleted U-residues are shown as asterisks. Canonical Watson-Crick base pairs (|); G:U base pairs (:). Previously identified start codons are doubled underlined. Potential upstream AUG start codons are indicated by wave underlines. Alternatively edited nucleotides are shown in red. Common anchor regions are shown in blue. *ND7* (D), and *RPS12* (E): Predicted mRNA and protein sequences, based on identified gRNAs.

The identification of gRNAs that could alter the reading frame led us to re-analyze the ORFs of the edited transcripts. In addition to *CR3*, we found extended ORFs in two different frames for Cytosine-rich Region 4 (*CR4*) and NADH Dehydrogenase (ND) subunit 9, while several others had shorter, but still significant ORFs in alternative frames (Fig 2). We do note that the original sequence publications for both *CR4* and *RPS12* (*CR6*) had indicated that the fully edited sequence contained extended ORFs in two different frames [26, 27]. Additionally, NADH Dehydrogenase subunit 3 (*ND3*) was also considered to be potentially dual coding, based on mutational analysis described below.

**Fig 2.**
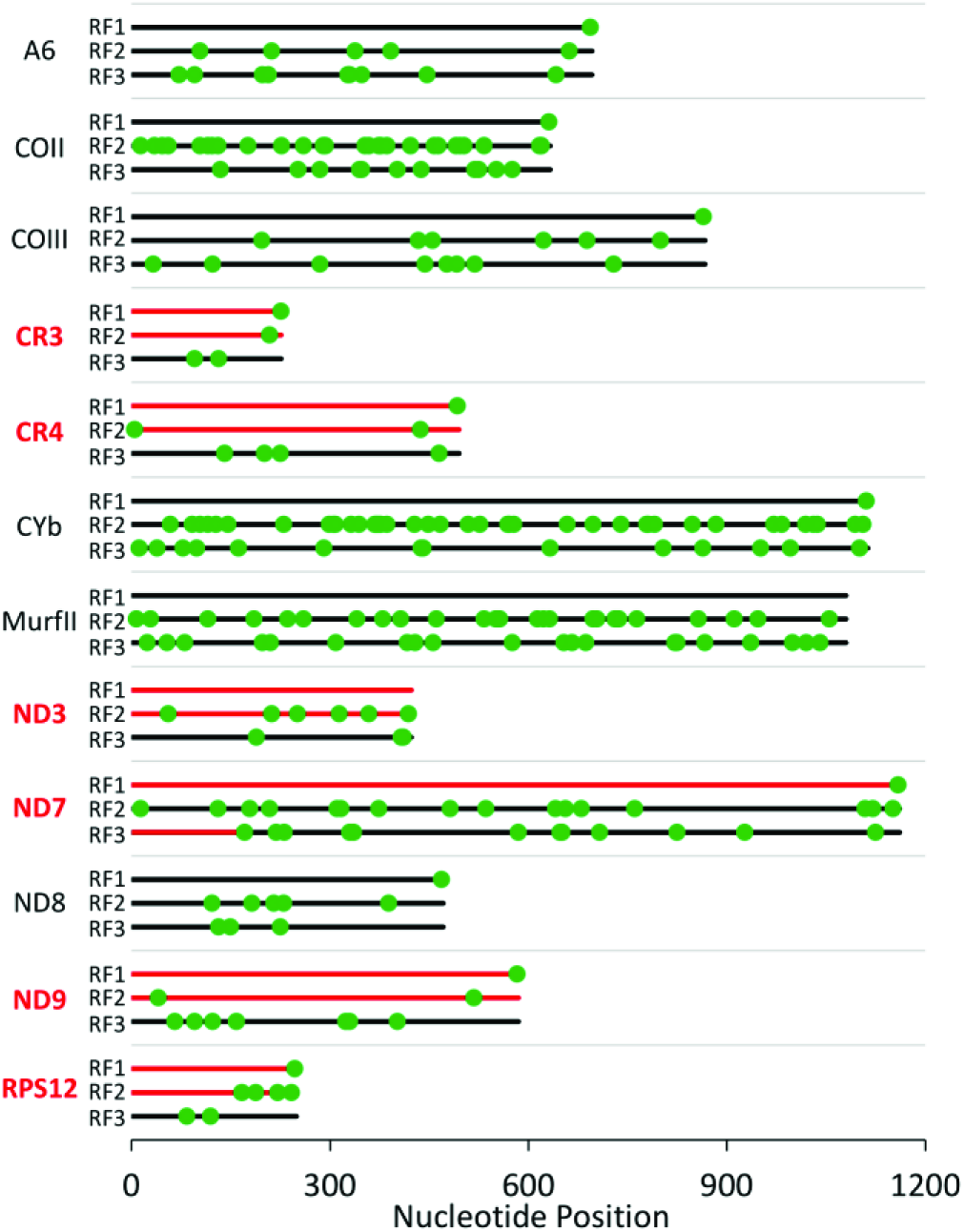
Positions of stop codons on all RFs of the edited genes in *T. brucei*. For each gene, reading frame 1 (RF1) is designated as the protein ORF previously identified. Hypothetical dual-coding reading frames are shown in red. *A6* = ATPase 6; *CO* = Cytochrome Oxidase; *CYb* = Cytochrome b; *Murf* = Maxicircle unidentified reading frame; *ND3* - *ND9* = NADH Dehydrogenase subunits.

As we did find potential ARFs in the edited transcripts, we analyzed the predicted ORFs for biases in their mutational pattern. Dual-coding genes often display an atypical codon mutation bias due to constraints imposed by the need to maintain protein function in both genes. In single-coding genes, changes in the third nucleotide of a codon give rise to synonymous amino acids, so this position (N3) is much less constrained. In contrast, in dual-coding genes, the N3 position in one frame is the N1 or N2 position in the alternative frame. Therefore, they have low rates of synonymous mutations [28]. This codon bias has been used to develop algorithms to detect novel overlapping genes [29, 30]. These algorithms however, cannot be used in the analysis of our edited transcripts as the two-component genetic system (mRNAs created by gRNA editing) introduces another layer of mutational constraint [31]. In addition, the edited sequence of the transcripts is known for only a limited number of kinetoplastids, and only the salivarian trypanosomes have the same general life cycle; other kinetoplastids, like *Leishmania* and *T. cruzi*, have evolved different infective cycles and are under very different selective pressures [5, 32, 33]. Fully edited sequences are known for *T. vivax*, the earliest branching salivarian trypanosome [34, 35]. *T. vivax* differs from *T. brucei* in that they complete the insect phase of their life cycle entirely within the proboscis of the fly. This parasite has been described as an intermediate stage in the evolutionary pathway from mechanical transmission (ancestral) to full adaptation to the midgut and salivary glands of the tsetse fly [36]. Using the *T. vivax* sequence, we analyzed mutation patterns in all of the mitochondrially-encoded mRNAs (Fig 3). mRNA sequences were aligned by codons based on their protein alignments (Clustal Omega [21]). Mutated codons were identified and classified as silent, missense and nonsense mutations. Missense mutations were further divided into three groups based on the PAM 250 matrix [22]. These data clearly show that the RNA editing process significantly constrains the types of mutations tolerated within the mitochondrial genome. In comparison to the genes that are not edited (*ND1*, *ND2*, *ND4*, *ND5*, *COI*) or have limited editing (*CYb*, *Murf II* and *COII*), a distinct suppression of silent mutations and strongly conserved missense mutations were observed for all of the pan-edited genes, consistent with previous observations (Fig 3) [31]. A suppression of mutations that lead to moderately conserved amino acid replacements was also observed, but these were not as striking due to the low frequency of this type of mutation. No significant difference was observed in the frequency of not conserved missense mutations, though a trend towards a lower frequency of these mutations in the putative dual-coding genes (*CR3*, *CR4*, *ND3*, *ND9*, *5’ND7* and *RPS12*) was noted. This was complemented by a significant increase in the frequency of strongly conserved missense mutations in the putative dual-coding genes in comparison to the other pan-edited genes (*3’ND7*, *ND8*, *A6* and *COIII*).

**Fig 3.**
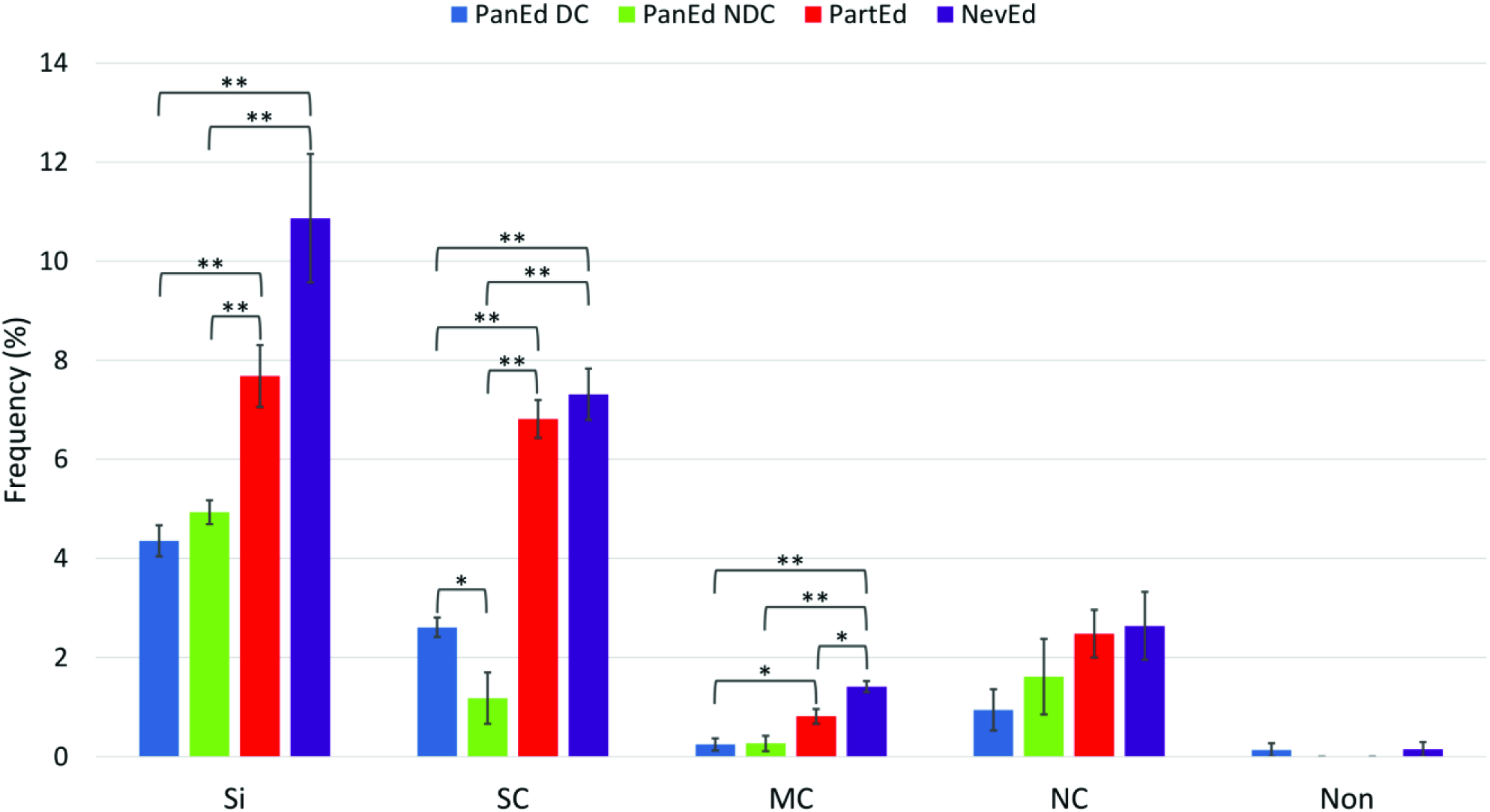
Mutational frequencies in mitochondrially encoded genes categorized by effect on amino acid sequence. *T. brucei* and *T. vivax* pan-edited dual-coding (PanEd DC), pan-edited nondual-coding (PanEd NDC), partially edited (PartEd), and never edited (NevEd) mRNA sequences were aligned based on their amino acid alignment (reading frame 1, defined as the reading frame encoding the gene product previously annotated in the literature) [21]. Mutations were categorized as silent (Si), strongly conserved (SC), modestly conserved (MC), not conserved (NC) or nonsense (Non). The amount of conservation was determined using the PAM 250 matrix, where conversions with a value 0 <x≤0.5 were considered modestly conserved, and conversions with a value >0.5 were considered strongly conserved, and conversions with a value ≤0 were considered not conserved. Error bars depict standard error. * p <0.05, ** p <0.01 (unpaired t-test).

Surprisingly, while the overall mutational frequency of the fully edited pan-edited genes was similar, a comparison of the conservation of editing patterns did show a significant difference between the putative dual-coding and the other pan-edited genes (Fig 4). The dual-coding genes consistently had a lower conservation of their editing pattern. Upon further examination, we found that most changes in the editing pattern resulted from thymidine insertions and deletions within the maxicircle DNA sequence, which was then corrected by the editing machinery. These types of mutations do not result in a change to the final mRNA sequence once edited. The *T. brucei* (*Tb*) dual-coding genes appeared to consistently insert more U-residues, while *T. vivax* (*Tv*) had more U-residues encoded within the DNA sequence. Indeed, comparisons of the length of the coding regions of *Tb* and *Tv* cryptogenes (unedited sequence) show that the putative dual-coding genes are almost 10% shorter in *Tb*. In contrast, the non-dual coding cryptogenes are not significantly shorter (~2.5%). Some of the other changes in editing patterns did generate small internal frameshifts as previously described by Landweber and Gilbert [31]. However, the high prevalence of internal frameshifts reported for *COIII* by Landweber and Gilbert is reflected in our analysis only for *COIII* and *A6*.

**Fig 4.**
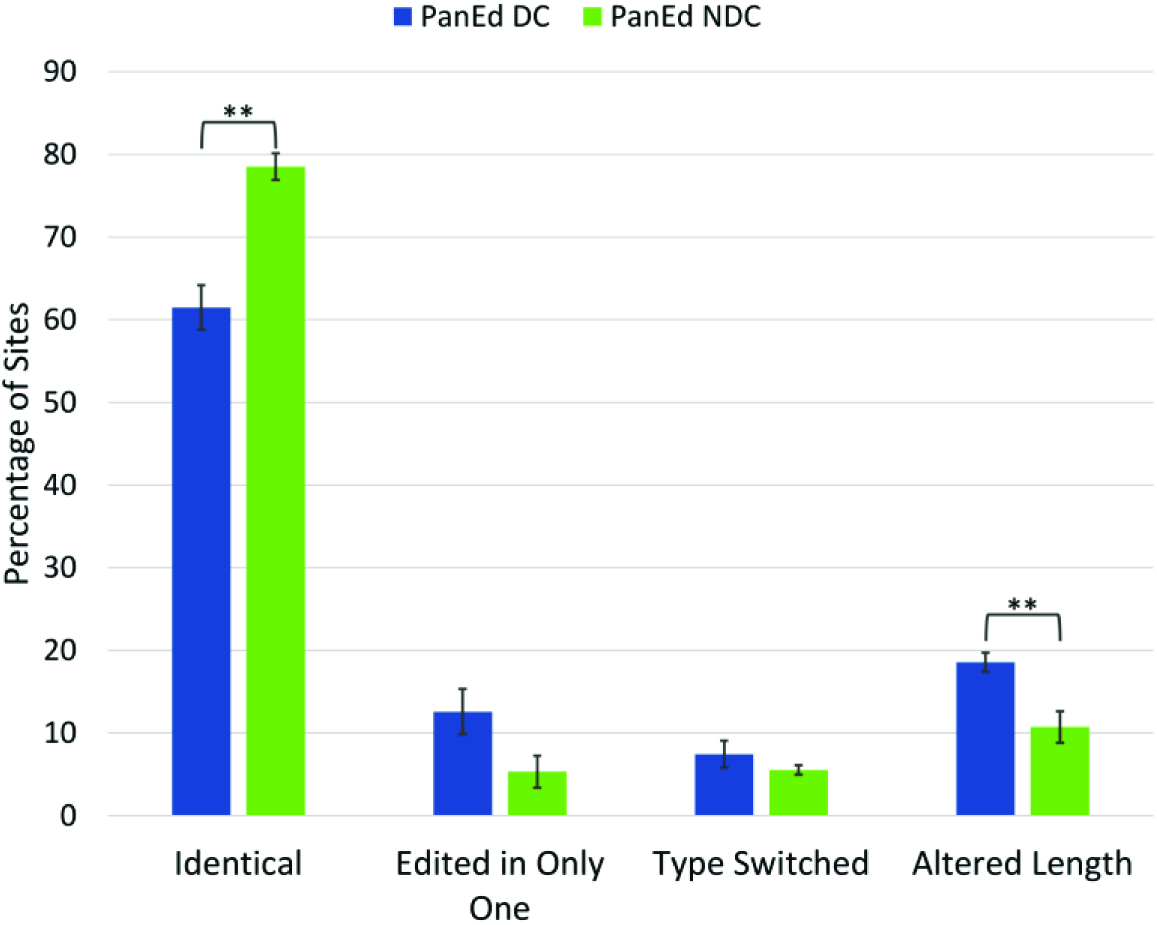
Percent Conservation of editing patterns between *T. brucei* and *T. vivax*. Alignment of the fully edited mRNAs was based on ACG sequence (see S1 Fig). Each editing site was defined as a site on at least one of the two aligned mRNAs where an editing event occurred. Sites were then classified as identical, only identified in one of the two sequences, type switched (one site is an insertion and the other is a deletion), or altered in length. Error bars depict standard error. * p <0.05, ** p < 0.01 (unpaired t-test).

Since differences in the types of amino acid mutations were observed, we performed a principal component analysis on the frequency of mutation types for all three reading frames of the pan-edited genes (Fig 5). In addition, we included the percentage of editing site conservation as a variable. This analysis clearly clustered the putative +1 dual-coding transcripts (reading frame 2). The first component (z-axis,) is strongly based on editing conservation, and separates the dual-coding genes from the other pan-edited genes as expected. While component 2 (x-axis) separated ORF1 and ORF3 from ORF2 of each gene, component 3 clearly separated the dual-coding ORF2s from nondual-coding ORF2s. The *ND7N* ORF3 was the only exception, and the gRNA data suggests that it is a dual-coding gene using the +2 (ORF3) reading frame. This suggests that an additional layer of mutational constraint beyond that imposed by the RNA editing process can be detected for six of the extensively edited transcripts.

**Fig 5.**
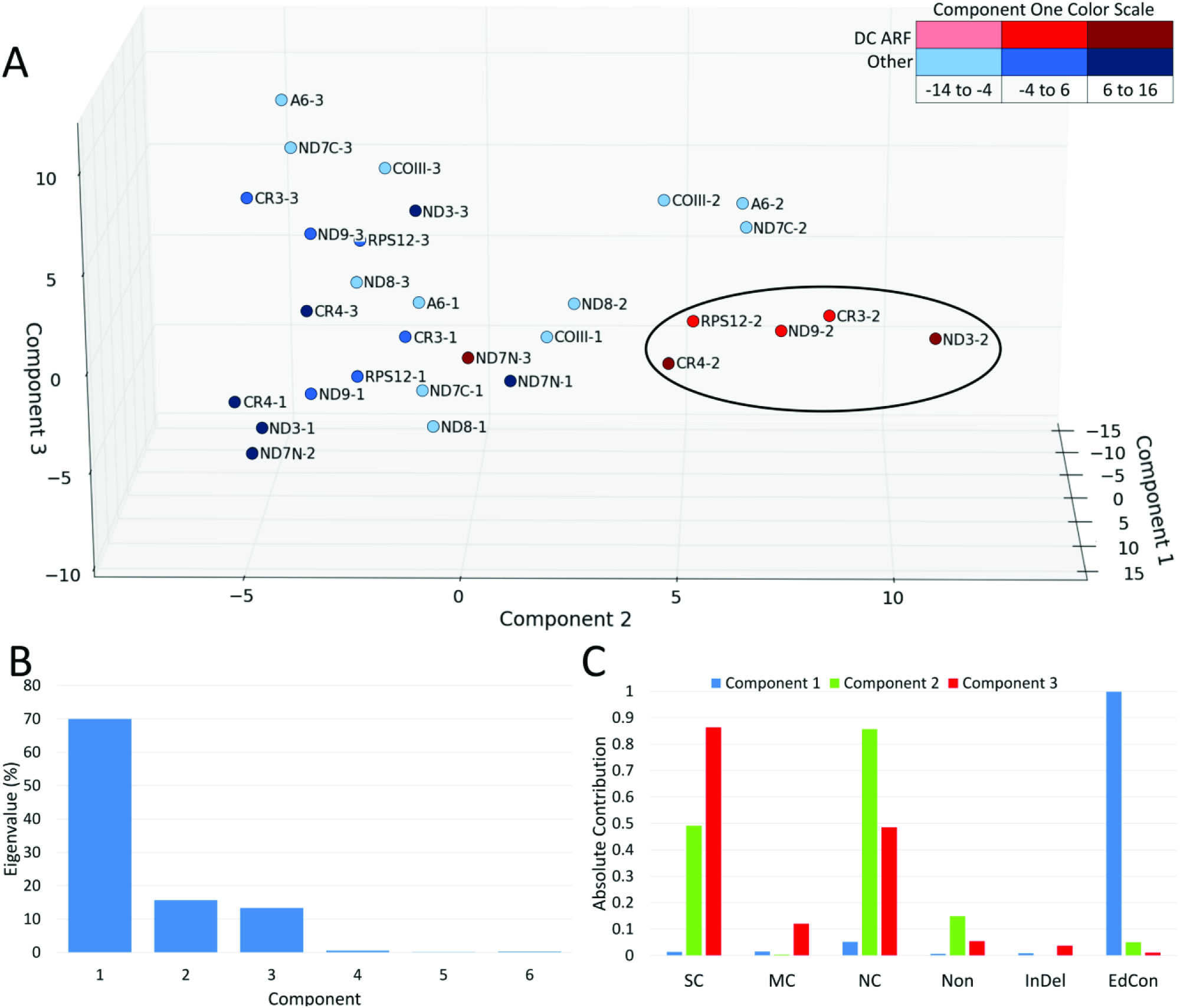
Principal component analysis of frequency of amino acid mutation types and editing conservation between *T. brucei* and *T. vivax* pan-edited transcripts. **A.** First factorial plan (z-axis: first component, x-axis: second component, y-axis: third component). ND7N = ND7 5’ editing domain, ND7C = ND7 3’ editing domain. ORF2 and ORF3 are defined as the +1 and +2 reading frames, respectively. **B**. Histogram of eigen values for first six components. Eigen values represent the amount of the variance accounted for by each component. **C**. Absolute contribution of each analyzed mutation frequency to components 1, 2, and 3. Amount of conservation was determined using the PAM 250 matrix as described in Fig 3. Mutation type: SC = strongly conserved, MC = moderately conserved, NC = Not conserved, Non = Nonsense, InDel = insertion or deletion. Editing conservation (EdCon) was determined using alignments of edited mRNAs (S1 Fig). Aligned editing sites were characterized as identical or altered.

Because dual-coding genes are often conserved in multiple species, we analyzed the available sequences of other kinetoplastids (*Leishmania tarentolae (Lt), Leishmania mexicana amazonensis (Lma), Phytomonas serpens (Ps*)) to determine if they also contain multiple overlapping reading frames with homology to those found in *T. brucei*. Interestingly, many of the alternative reading frames did show some homology to the ARFs found in *Tb*. However, most of these ARFs are punctuated with stop codons (S2 Fig). Extended alternative reading frames are found in *CR3*, 5’*ND7* and *RPS12* in *Ps*. However, the extended ARF in the *Ps CR3* is in the +2 reading frame and the *ND7* and *RPS12* ARFs shows very little homology with the *Tb/Tv* ARF (S2 Fig A, D, F) [37]. The *L. tarentolae CR4* orthologue also has two extended ORFs. Interestingly, the published sequence for *Lt CR4* appears to switch between the two ORFs (switch appears to occur in a stretch of 13 inserted Us) [38]. This may explain why only the carboxyl half of the published *Lt CR4* showed good homology with *Tb* and *Lma* [39]. Translation of the *Lt* ARF does generate a protein with the N-terminus showing high homology to the conventional *Tb* and *Lma CR4*, while translation of the published ORF shows some homology to the *Tb CR4* ARF (S2 Fig B). These data are intriguing enough that these sequences should be re-examined. While most of the other pan-edited transcripts had multiple stop codons in the +1 and +2 reading frames, many did show good homology to the *Tb* ARF sequences. Particularly intriguing are the *ND3*, *ND8* and *ND9* alignments. While internal stop codons are found in *Tv*, *Lt* and *Lma ND9* ARFs, they show strong homology to the *Tb ND9* ARF throughout the protein (S2 Fig E). In *ND3*, the amino ends of the ARFs show strong homology between all four of the *Trypanosoma* and *Leishmania* species (S2 Fig C). This homology decreases after an internal stop codon found in the same position in 3 of the 4 species. As *ND8* is the only other pan-edited gene in *Lt*, *Lam* and *Ps*, we also examined the conservation of the *ND8* ORF and ARF, even though our mutational analyses did not tag the *ND8* gene as dual coding. While the *ND8* ARFs were punctuated by multiple stop codons, they surprisingly also showed areas of strong homology between all 5 species, especially down stream of an internal methionine (S2 Fig G). We do note that we cannot rule out the possibility that alternative editing can remove stop codons observed in the ARFs.

Analyses of the ARF predicted proteins suggest that they are all short transmembrane proteins with two or more predicted transmembrane alpha helices (Fig 6) [40, 41]. While functional homologues are often difficult to detect in trypanosomes, searches using the predicted protein sequence of each ARF did identify small molecule transport proteins with limited confidence. Using Phyre2, the *ND7* ARF was identified as a homolog of the bacterial sugar transporter SemiSWEET (61.5% confidence) [42, 43]. SemiSWEET, which forms homodimeric structures, is also a distant homolog of the yeast mitochondrial pyruvate carrier 1 (MPC1). This protein has two transmembrane alpha helices and forms a heterodimer with either of the other two pyruvate carrier proteins [44, 45]. While still very speculative, it is intriguing that the small ARF proteins might oligomerize to form small mitochondrial membrane transporters.

**Fig 6.**
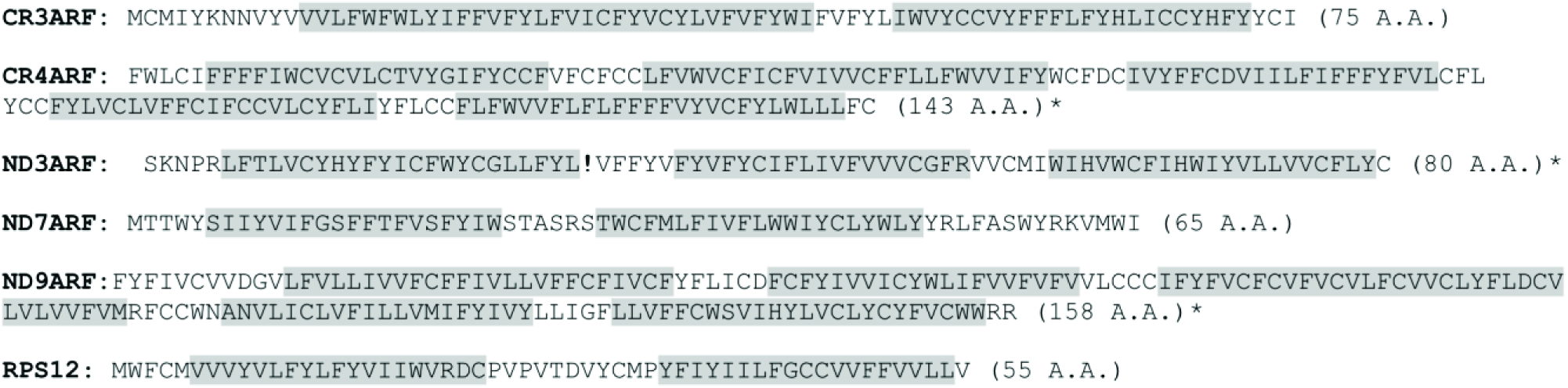
Amino acid sequences of ARFs of dual-coding genes. Predicted transmembrane regions are shaded in gray [40, 41]. *No start codon was identified and the amino acid sequence shown begins at the 5’ end or after the first stop codon at the 5’ end. Exclamation point indicates premature termination codon.

## Discussion

The work presented here, suggests that as many as six of the extensively edited mRNAs in *T. brucei* are dual-coding and that it is alternative editing using different terminal gRNAs that allows access to the two different reading frames. Deep sequencing of the 5’ end of *CR3* indicates fully edited transcripts that have access to both reading frames are present in the mitochondrial transcriptome and gRNA analyses indicate that three different cell lines contain gRNAs that can alternatively edit the 5’ ends of *CR3*, *RPS12* and *ND7*. In addition, analyses of the mutational bias in pan-edited genes suggest that an additional layer of mutational constraint is observed in the putative dual-coding genes. While the overall mutational frequency observed for the fully edited mRNAs is similar for all pan-edited genes, the types of amino acid changes that appear to be tolerated are significantly different. This is consistent with these genes having to maintain functional proteins in two different reading frames. Analyses of other trypanosomes, do show that some of the ARFs have intriguing homology to the ARFs identified in *T. brucei* and *T. vivax*. However, most of the ARFs are punctuated with stop codons. These data are difficult to interpret because we cannot rule out the possibility that the stop codons are removed by alternative editing events. In addition, the other trypanosome species have evolved very different infective life cycles and are under different selective pressures. For example, *P. serpens* is a pathogen that infects important crops and is transmitted by sap-feeding bugs. These parasites have glucose readily available in both life cycle stages and are unique in that they lack a fully functional respiratory electron transport chain [46, 47]. For *Leishmania*, all life cycle stages possess an active Krebs cycle and ETC linked to the generation of ATP [48, 49, 50]. These unique adaptations to different hosts suggest that they may not be under the same evolutionary pressure to maintain dual-coding genes.

Overlapping reading frames are common in viruses, and are thought to persist due to strong genome size constraints [51, 52]. More recently however, over-lapping genes have been identified in mammalian and bacterial genomes [53–56]. In these organisms, size is not an issue and the potential advantage of overlapping genes is less clear. For dual-coding genes, the need to maintain both ORFs constrains the ability of each protein to become optimally adapted [15]. As this constraint can be alleviated by gene duplication, it is thought that dual-coding regions can survive long evolutionary spans only if the overlap provides a selective advantage. In mammals, many of the identified dual-coding genes like *Gnas1* an*d XBP1*, produce two proteins that bind and regulate each other [57-58]. For these proteins, dual-coding may be advantageous for the tight co-expression needed. An alternative model, suggests that under high mutation rates, the overlapping of critical nucleotide residues is advantageous because it may reduce the target size for lethal mutations [59]. This may be particularly important for organisms that have evolved to exist in dual-metabolic environments (two hosts). We hypothesize that the trypanosome mitochondrial ARFs encode small metabolite transporters that provide a distinct growth advantage to bloodstream form parasites. The complete overlap of these small transporter genes with electron transport chain (ETC) genes would protect the integrity of the ETC genes that are required only in the insect host. Thus, in trypanosomes, dual-coding genes may be a mechanism to combat genetic drift during extended periods of growth in non-selective environments. In *T. brucei*, it is known that a number of bloodstream form essential proteins are functionally linked to Krebs cycle or ETC genes. While not a “classic” dual-coding gene in that production of the alternative protein does not involve overlapping reading frames, the pan-edited *COIII* gene does contain the information for two distinct proteins, COIII and AEP-1. AEP-1 is important for kinetoplastid DNA maintenance and over-expression of the DNA-binding domain results in a dominant negative phenotype including decreased cell growth and aberrant mitochondrial DNA structure [12]. The nuclear encoded α-ketoglutarate dehydrogenase E2 (α-KDE2) is known to be a dual-function protein, in that it plays important roles in both the Krebs cycle and in mitochondrial DNA inheritance [60]. RNAi knockdowns of this gene in bloodstream form (BF) trypanosomes also show a pronounced reduction in cell growth. Similarly, the Krebs cycle enzyme α-ketoglutarate decarboxylase (α-KDE1) is also a dual-function protein with overlapping targeting signals that allow it to be localized to both the mitochondrion and glycosomes [61]. RNAi knockdowns of α-KDE1 in BF trypanosomes is lethal, suggesting that in addition to its enzymatic role in the Krebs cycle, it plays an essential role in glycosomal function in *T. brucei* [61]. It has been previously suggested that both alternative editing and dual-function proteins are important mechanisms for expanding the functional diversity of proteins found in trypanosomes [11, 60-62]. We hypothesize, that in salivarian trypanosomes, an equally important role for these dual-coding/function genes may be the protection of genetic information.

The “why” of the unique RNA editing process in kinetoplastids has been a long-standing paradox. The complex machinery and the sheer number of gRNAs required to direct the thousands of U-insertion/deletions indicate that this process is metabolically very costly. Initially, it was proposed that U-insertion/deletion editing (kRNA editing) was one of many RNA editing processes that were in fact relics of the RNA world. However, the very different mechanism of the RNA editing systems in existence, and their very limited distribution within specific groups of organisms indicate that they are more likely derived traits that evolved later in evolution [63, 64]. The sheer complexity of the kRNA editing process, with no obvious selective advantage, led to the proposal that insertion/deletion editing arose via a constructive neutral evolution (CNE) pathway [65]. Indeed, RNA editing in trypanosomes is always mentioned in support of CNE as an example of how seemingly non-advantageous, complex processes can arise [66, 67]. More recently however, it has been hypothesized that RNA editing co-evolved with G-quadruplex structures found in the pre-edited mRNAs [16]. These structures are thought to be advantageous in that they can help regulate transcription in order to promote DNA replication and prevent kinetoplast DNA loss. However, they must be removed by the RNA editing system prior to translation [16]. Another prominent hypothesis is that RNA editing is advantageous because it is a mechanism by which an organism can fragment and scatter essential genetic information throughout a genome [6, 68]. Kinetoplast DNA is far less stable than chromosomal DNA, and loss of minicircles due to asymmetric division of the kDNA network have been frequently observed, especially in laboratory cultures of *Leishmania* [38, 69]. Buhrman et al. [68] suggest that the scattering of essential guide RNA genes throughout the DNA network, would prevent fast growing deletion mutants from outcompeting more metabolically versatile parasites during growth in the mammalian host. Using a mathematical model of gene fragmentation in changing environments (absence of functional selection), they showed a distinct advantage for gene fragmentation. In their model, the number of tolerable generations under periods of relaxed selective pressure was increased by more than 40% before loss of the ability to move to the next life cycle stage. If the dual-coding ARFs give BF trypanosomes a selective growth advantage similar to that observed by the *COIII* alternative protein AEP1, then the number of “essential” gRNA genes would increase greatly. Currently, only AEP1, *A6* and *RPS12* mitochondrial genes have been experimentally shown to be essential [12, 70-71]. In addition, the presence of alternative editing and dual-coding genes would complement the protection provided by gene fragmentation by also shielding the genes from deleterious point mutations within critical ETC genes. This suggests that the complex RNA editing system found in the mitochondria may therefore provide multiple molecular strategies to increase genetic robustness. Protection of the mitochondrial genome during growth in the mammal would increase the capacity for successful transfer to an insect vector and maximize the parasites long-term survival and spread.

## Acknowledgements

We thank the Ken Stuart Lab for trypanosome cell lines and Chris Adami for helpful discussions. We would also like to acknowledge the Berttina B. Wentworth Endowed Fund and the Dr. Marvis A. Richardson Endowed Fellowship Fund for their recognition of LEK.

## Supplementary Information

**S1 Fig.**
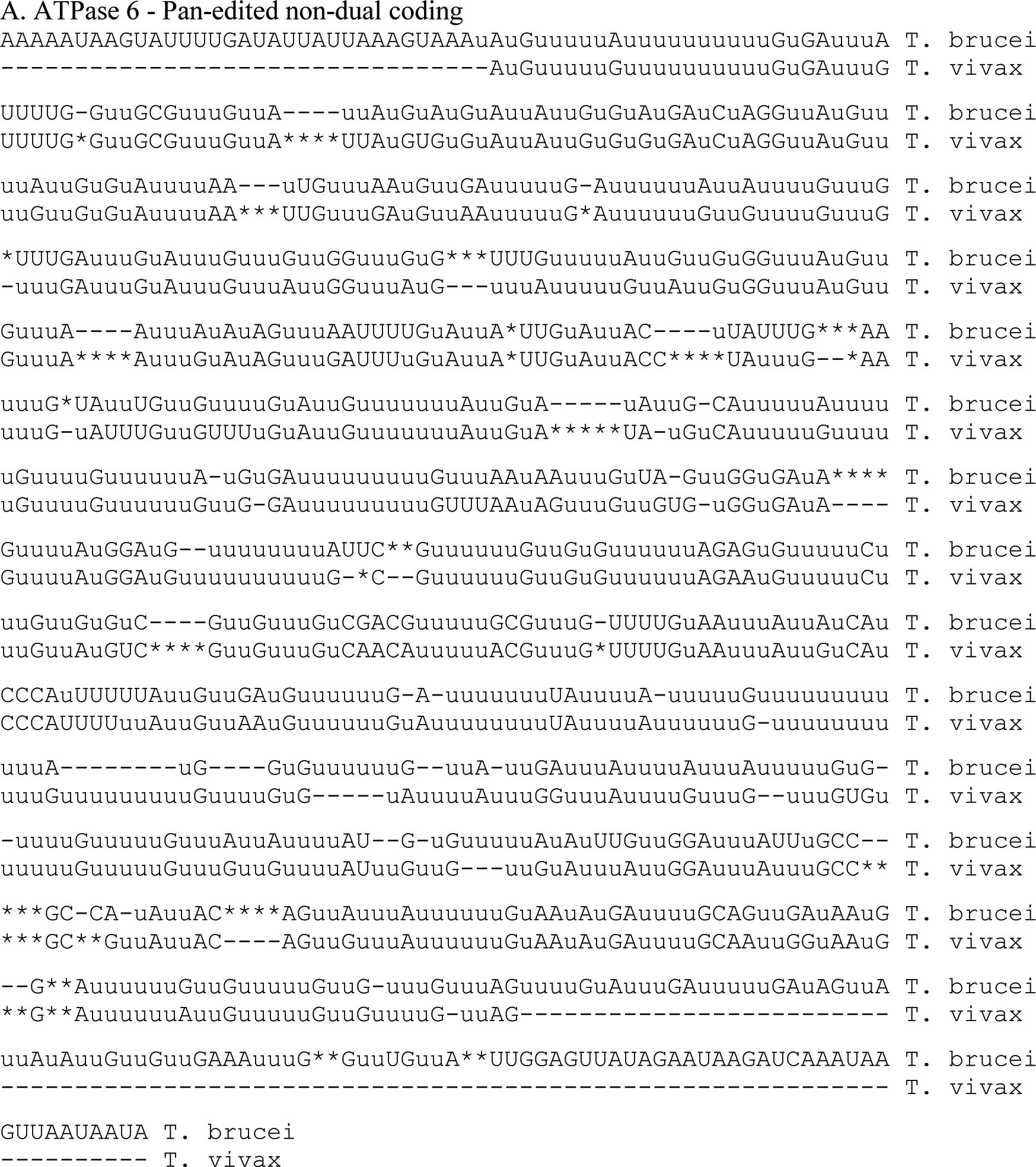
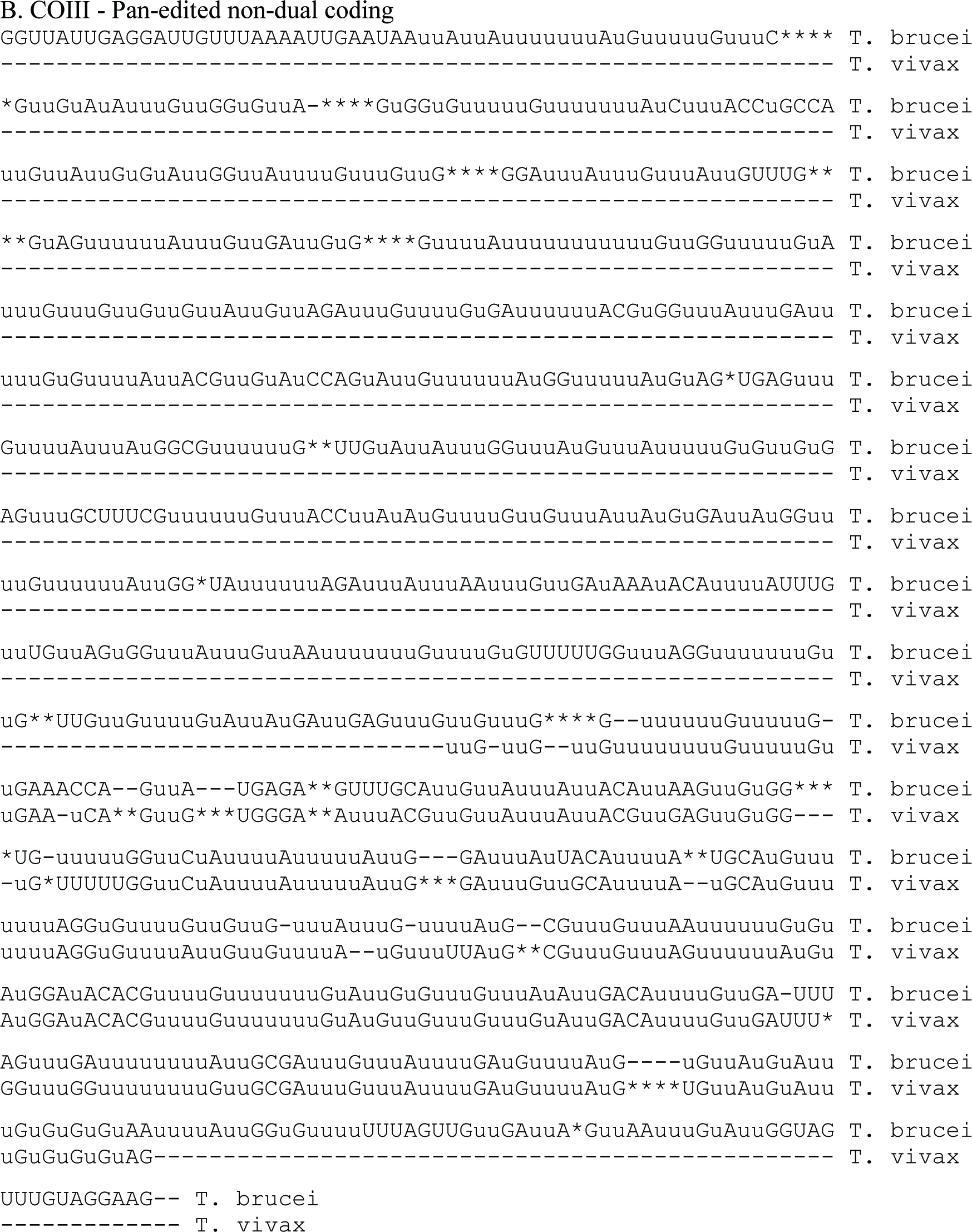
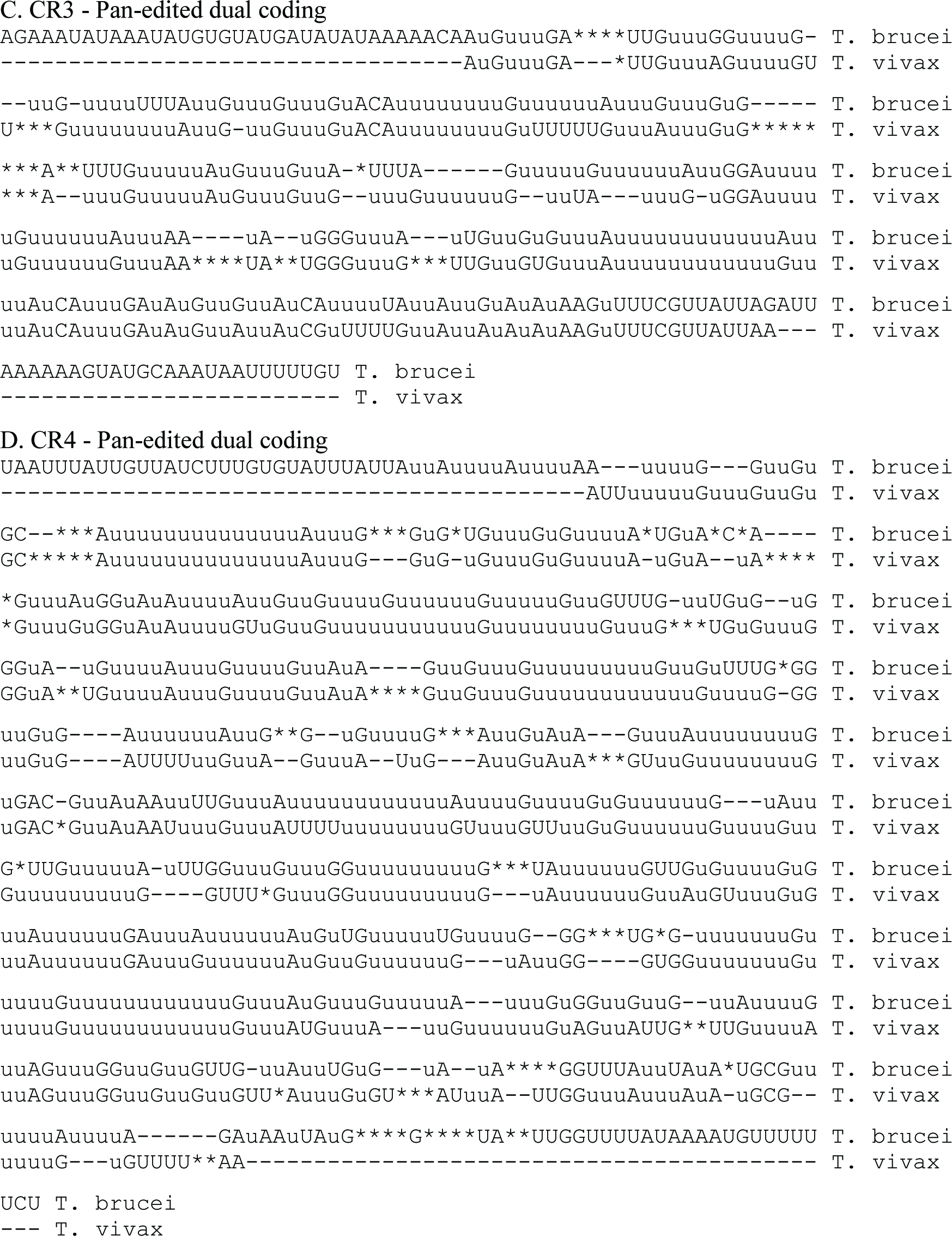
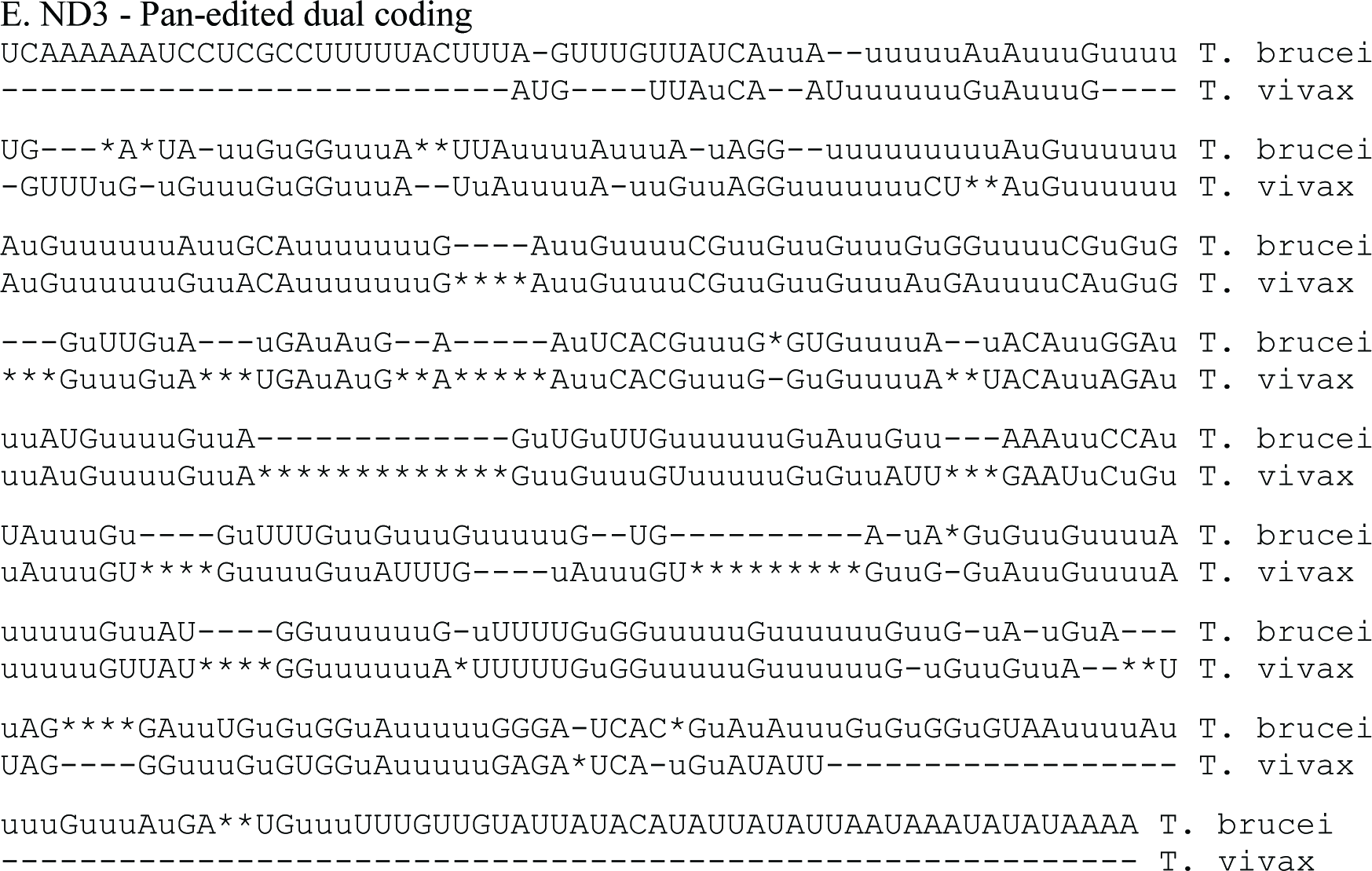
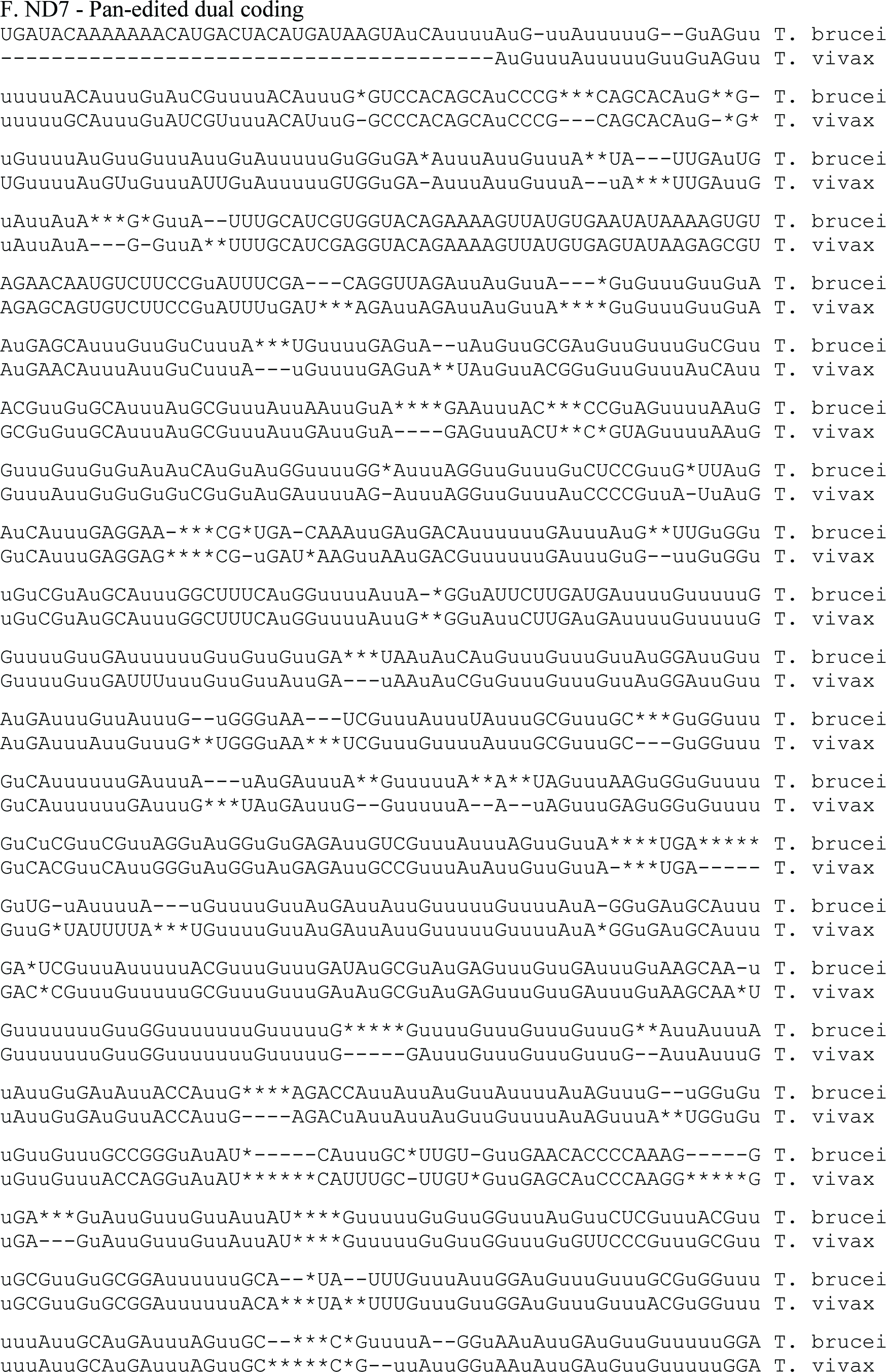
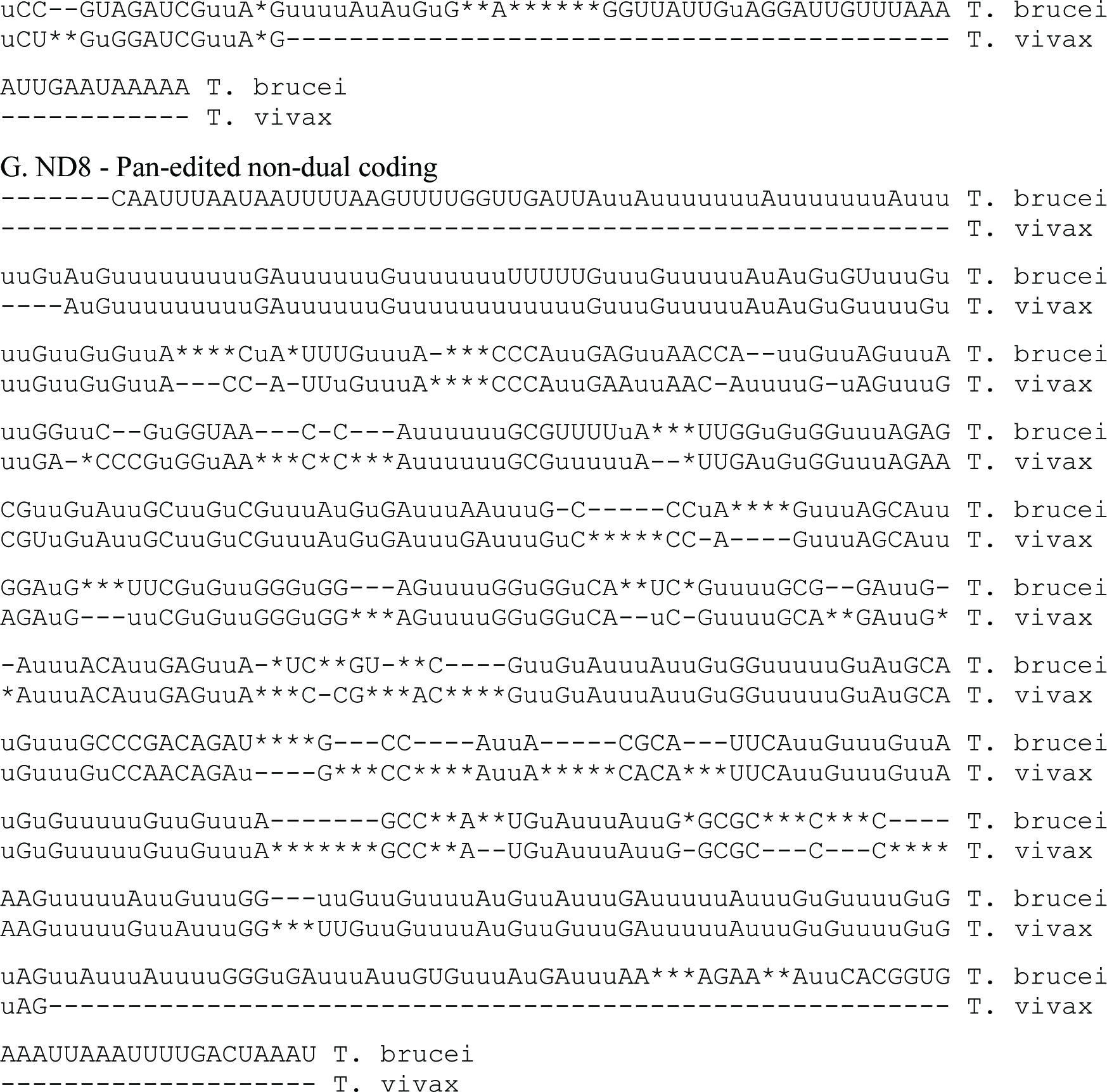
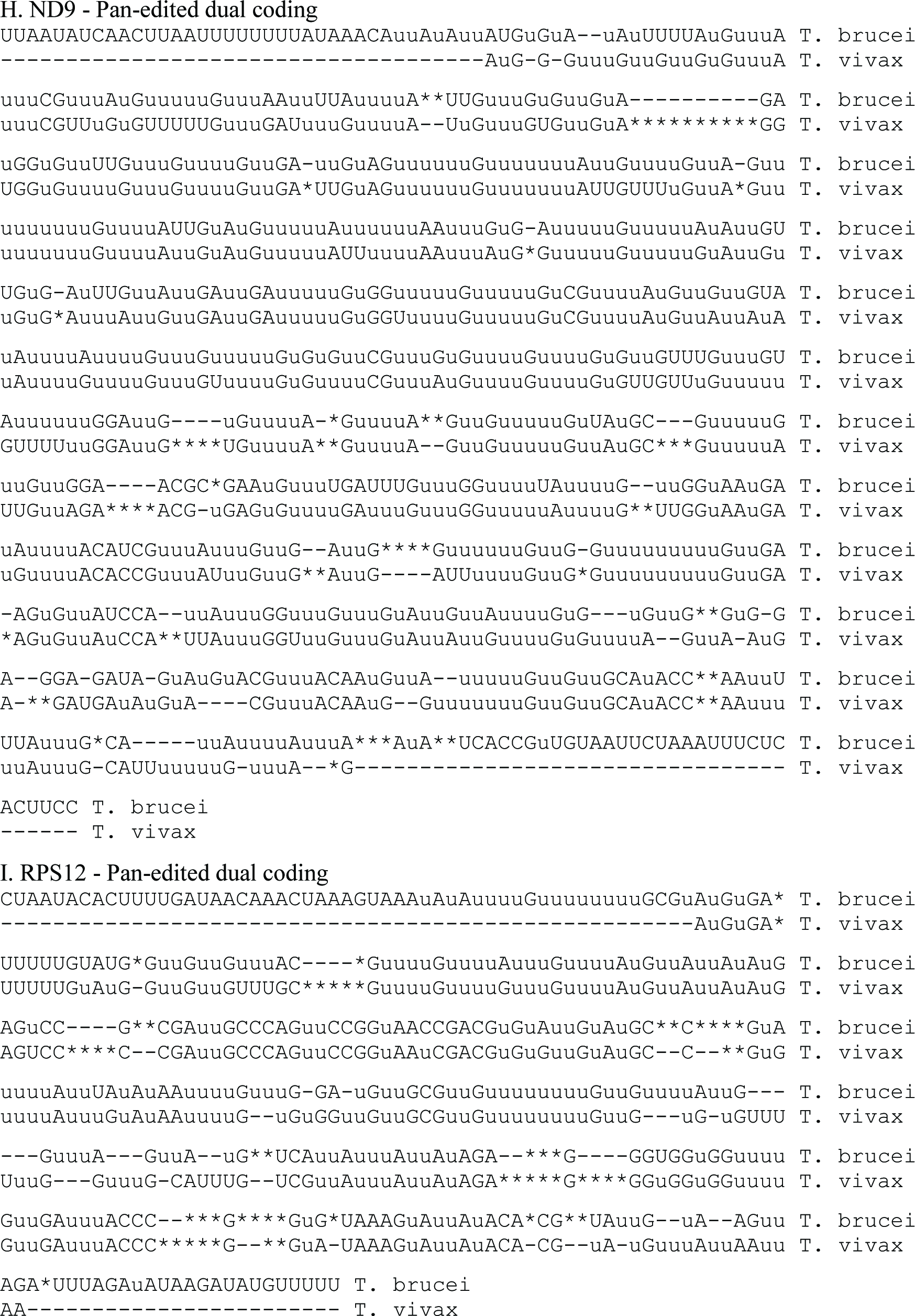
Alignments of *T. brucei* and *T. vivax* pan-edited mRNAs. *ATPase 6* (A), *COIII* (B), *CR3* (C), *CR4* (D), *ND3* (E), *ND7* (F), *ND8* (G), *ND9* (H), and *RPS12* (I). Uppercase letters indicate nucleotides originally encoded in the DNA, lower case u’s indicate uridines inserted during editing and asterisks indicate uridines removed during editing.

**S2 Fig.**
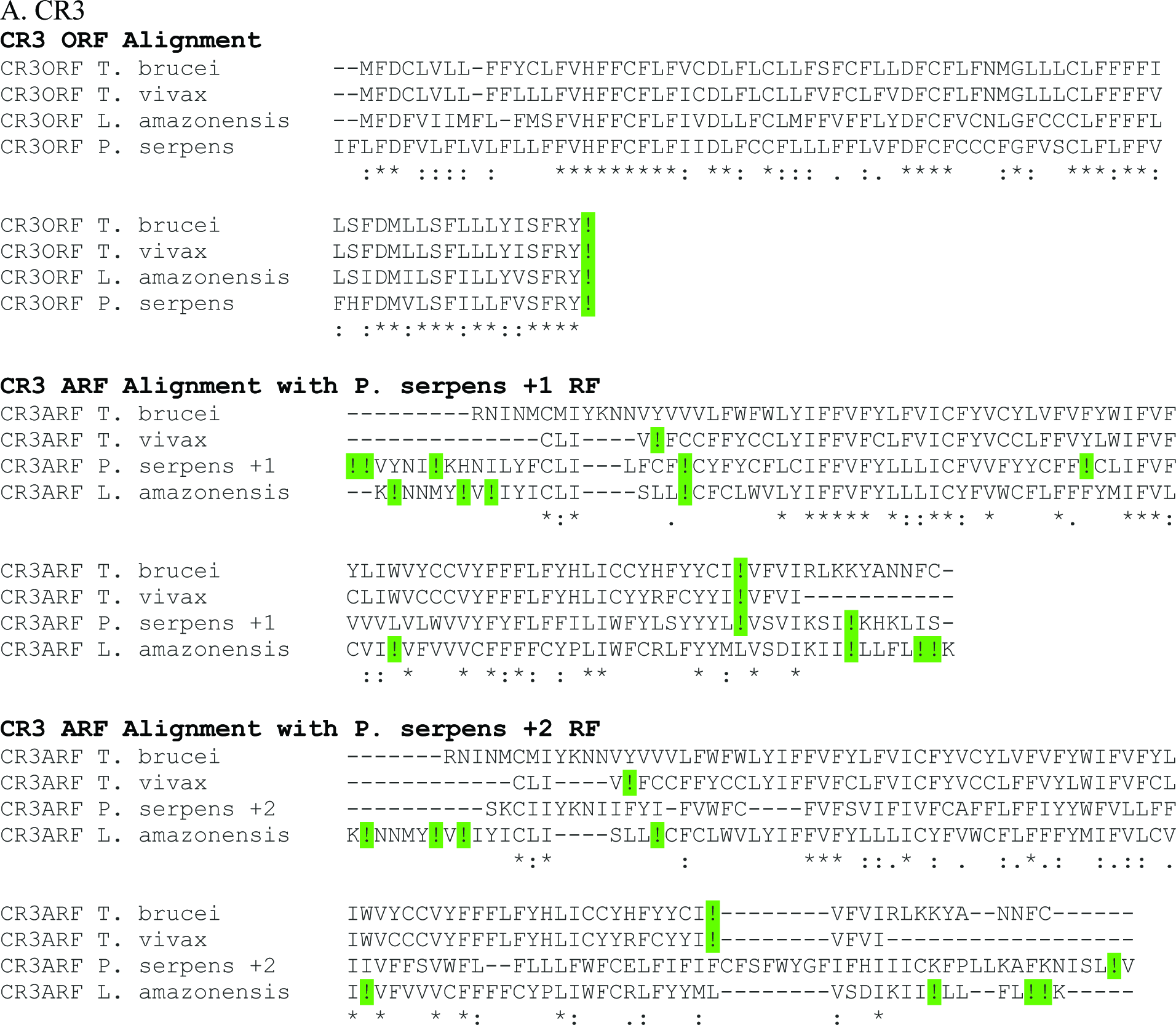
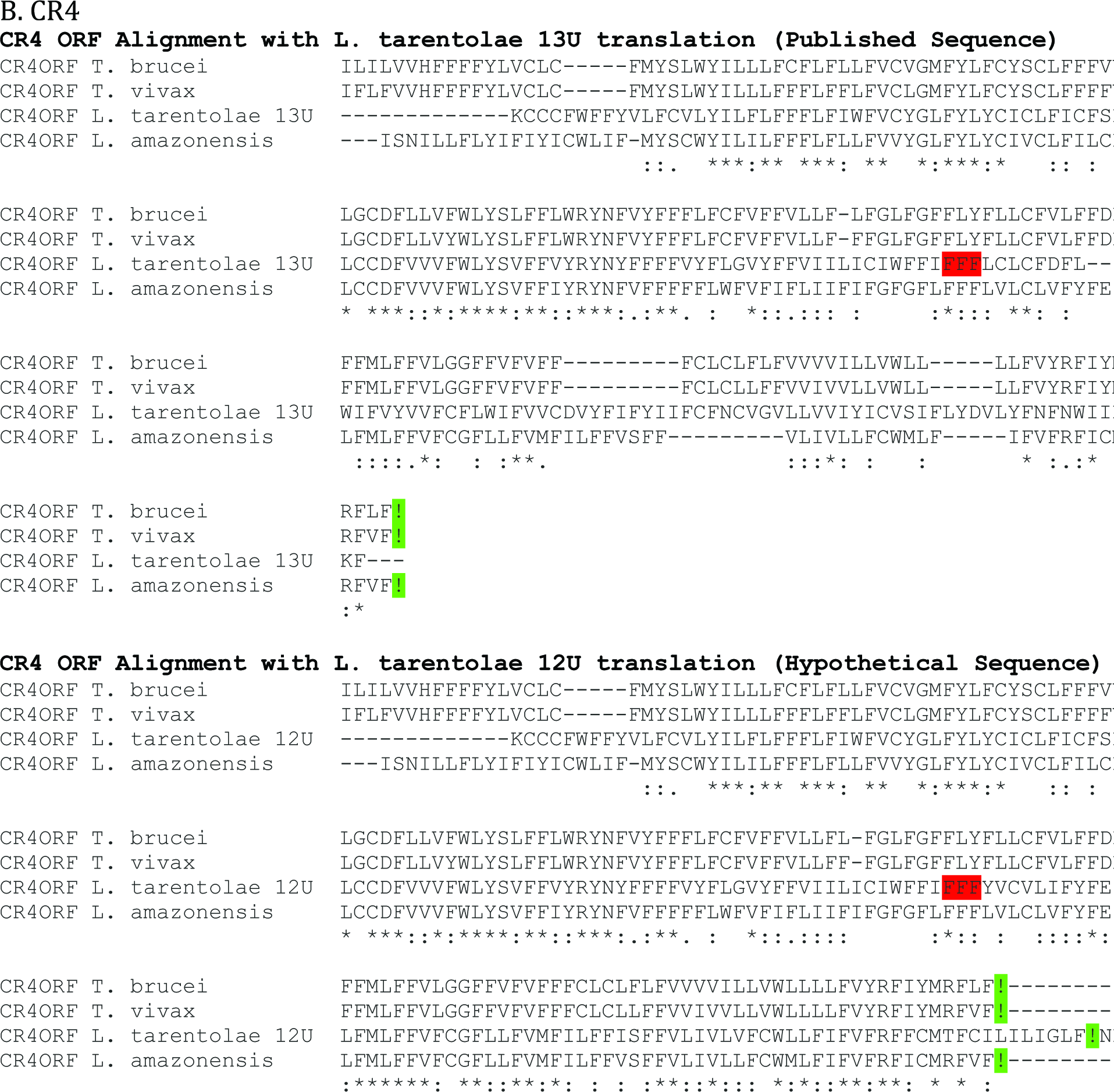
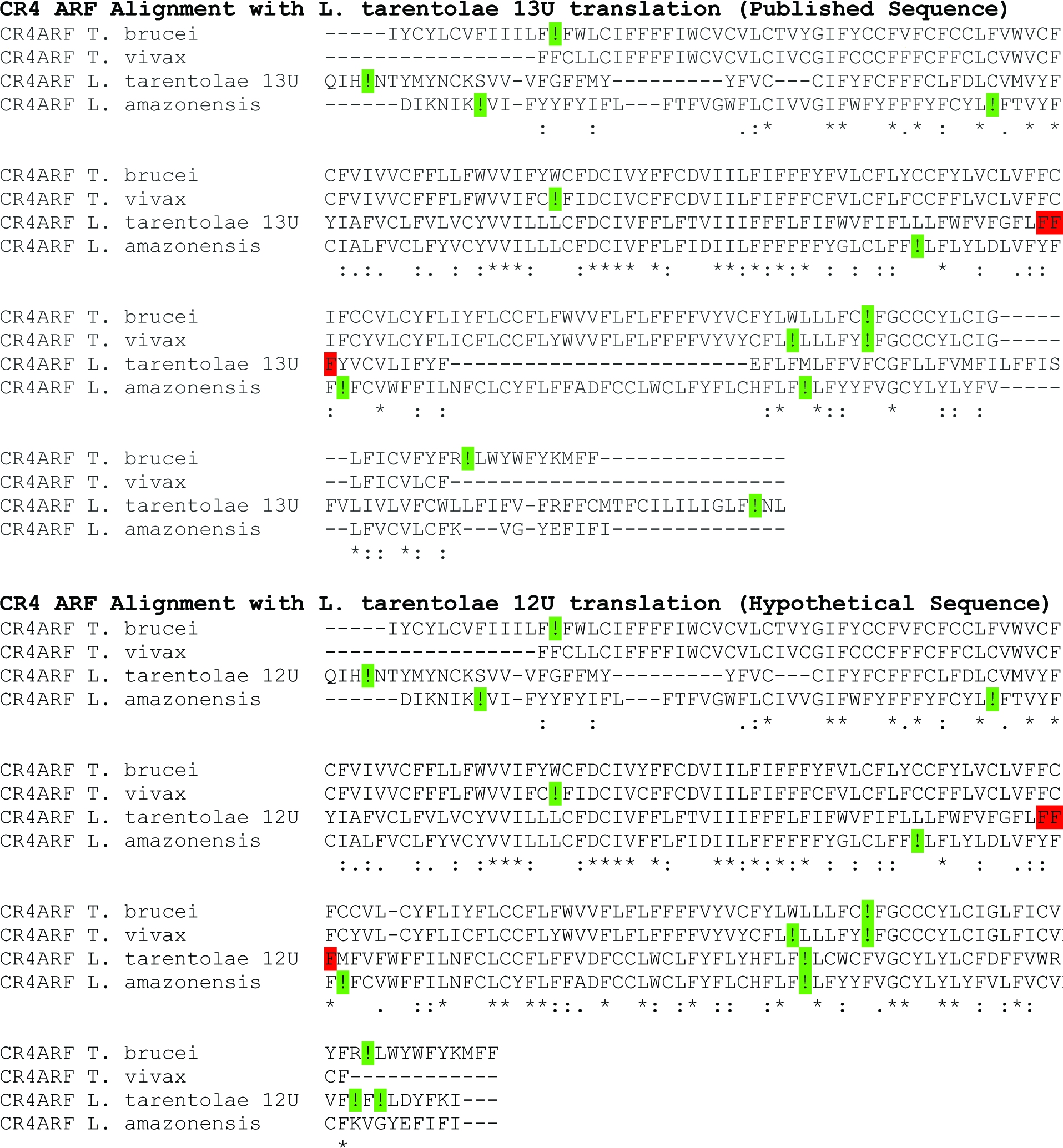
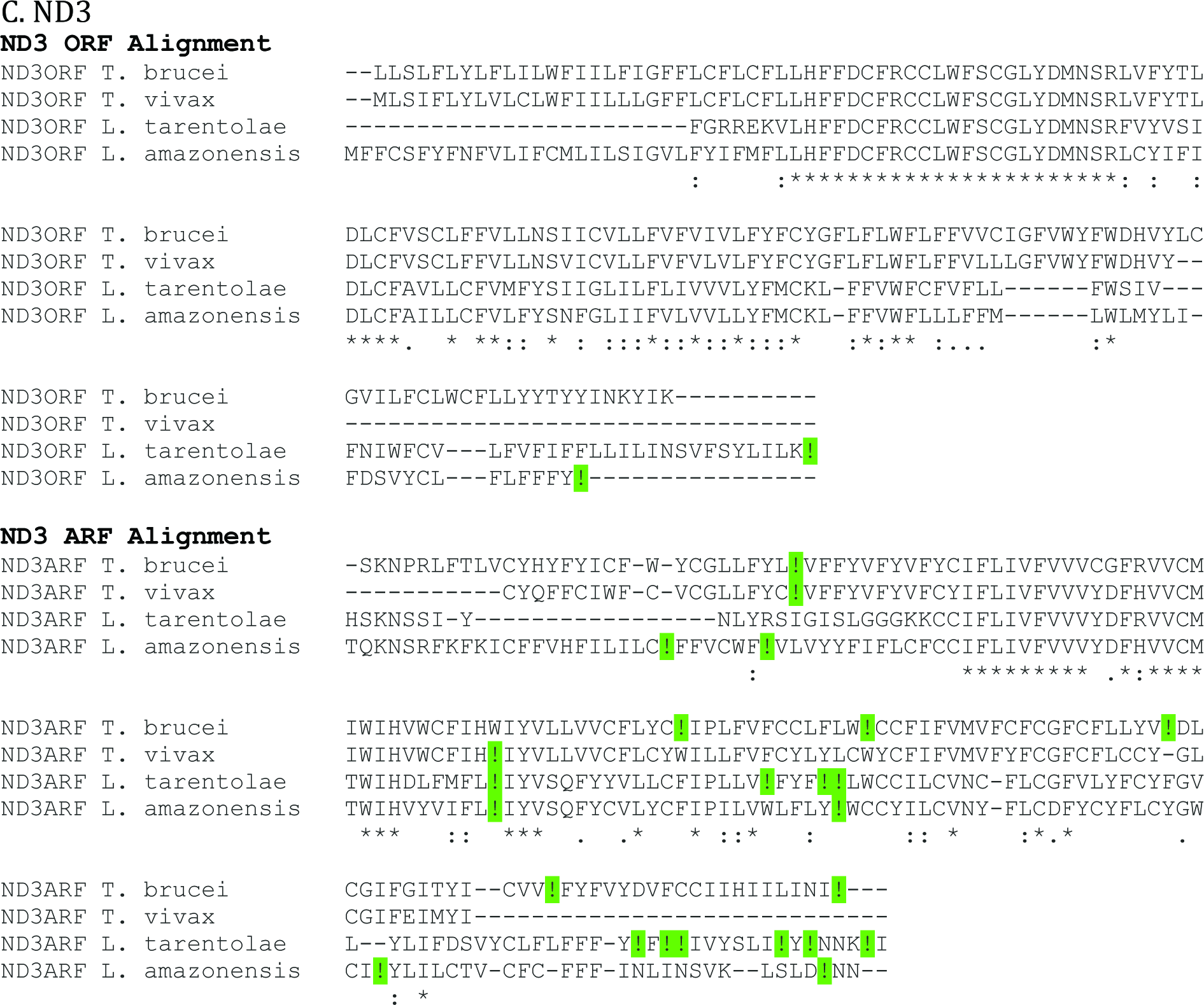
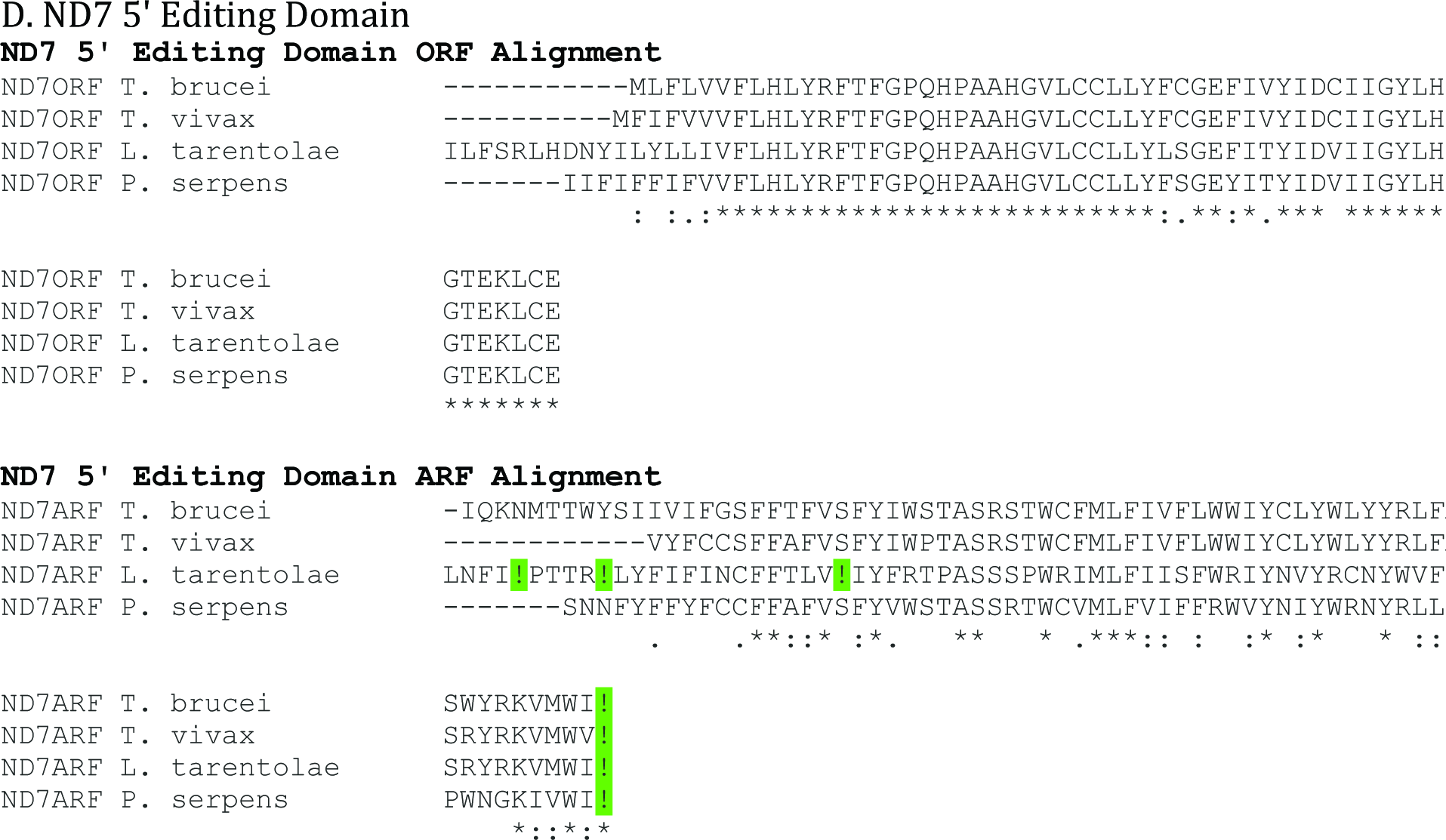
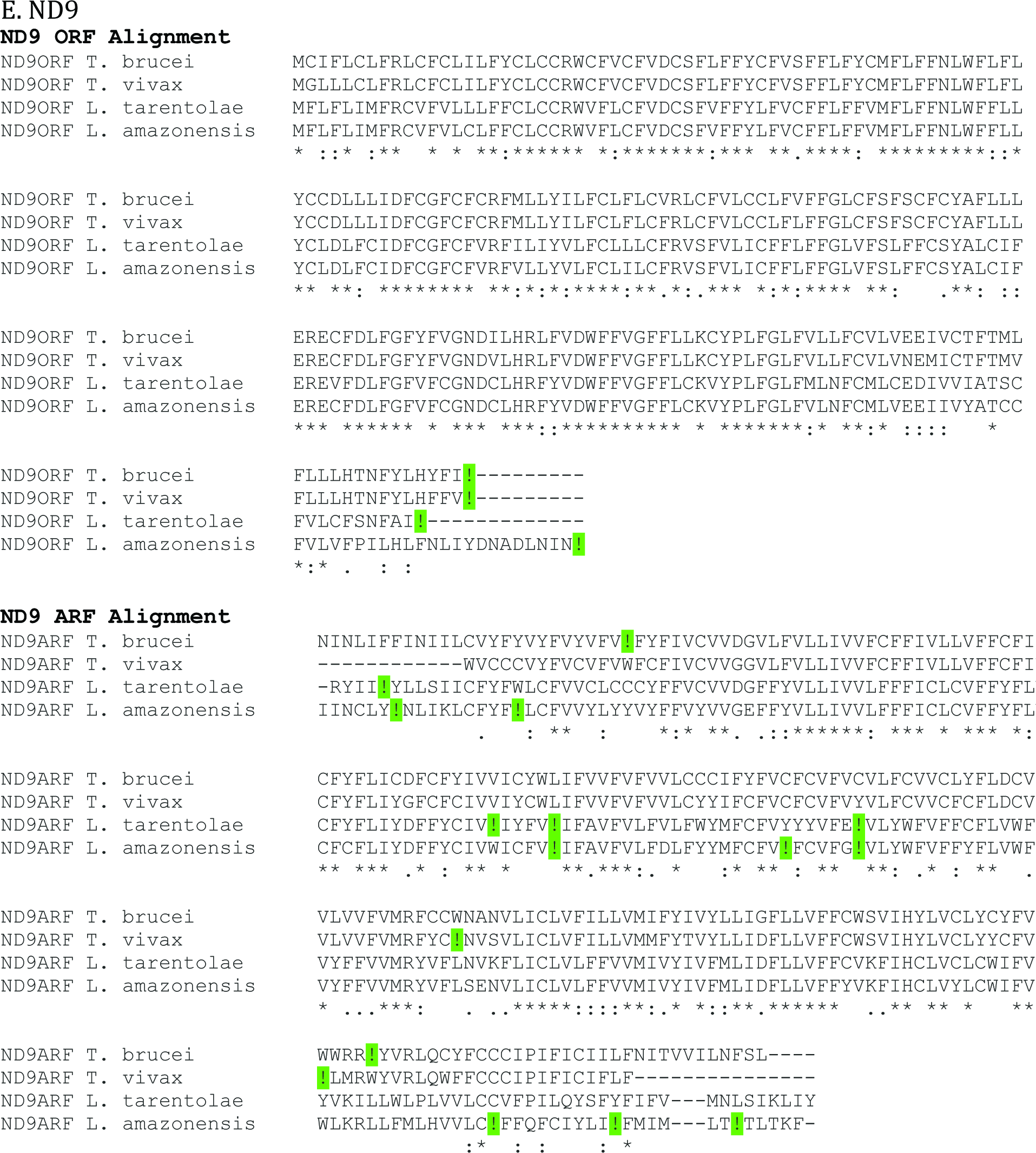
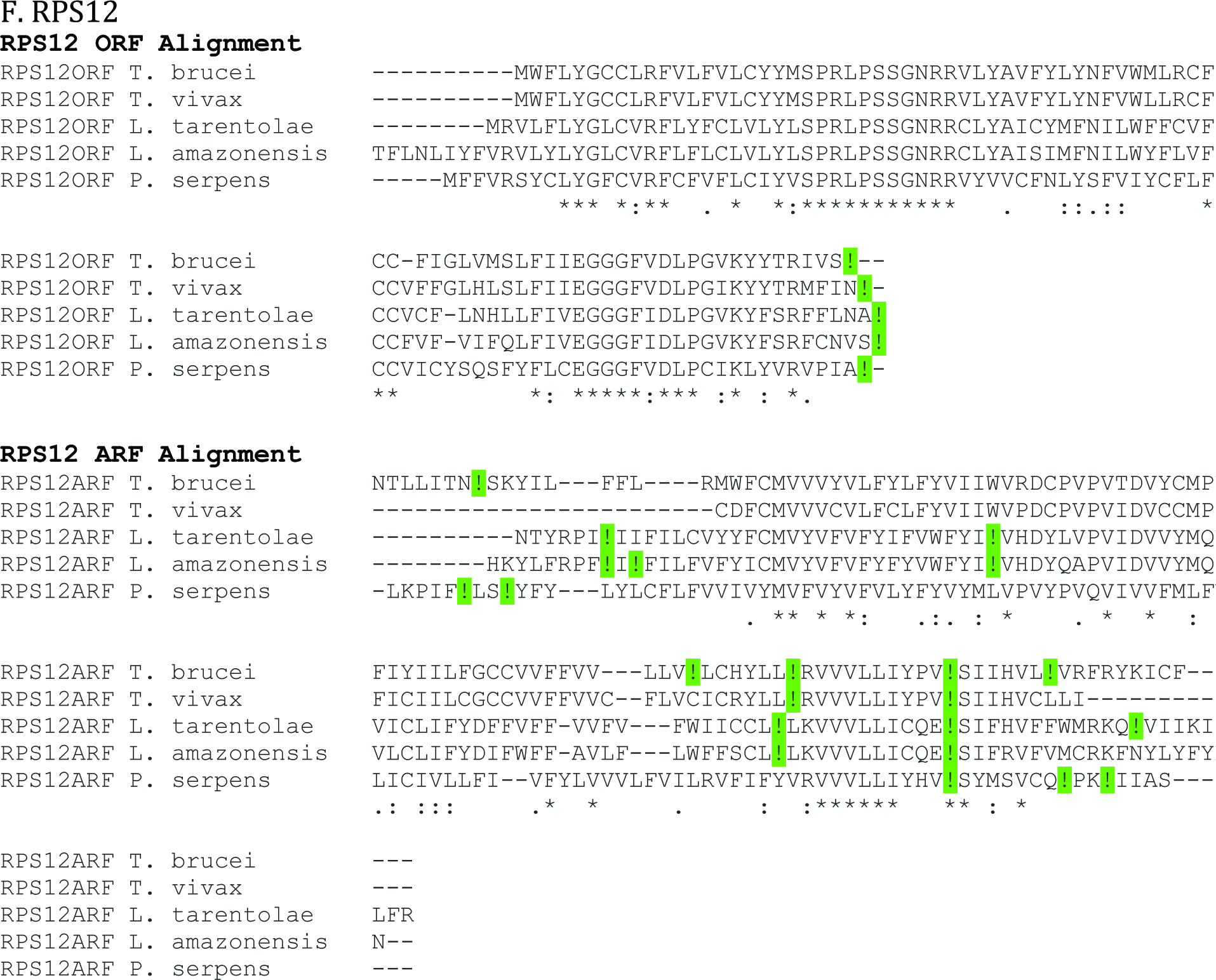
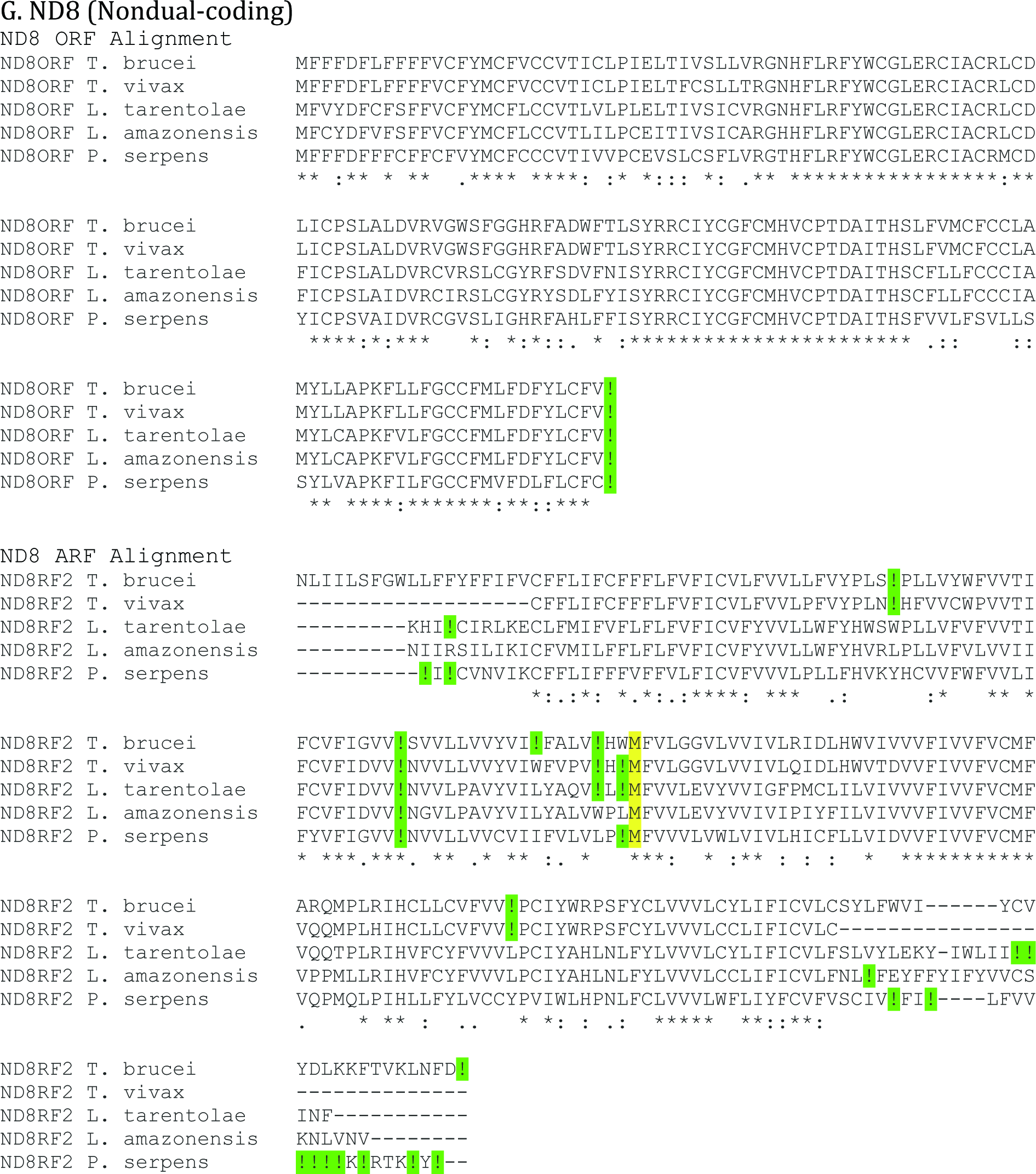
Alignments of protein sequences of pan-edited dual-coding genes in *L. tarentolae*, *L. amazonensis* and *P. serpens* with *T. brucei* and *T. vivax sequences*. A: *CR3*, B: *CR4*, C: *ND3*, D: *ND7* 5’ Editing Domain E: *ND9*, F: *RPS12*, G: *ND8* (Nondual-coding). Absent sequences were unavailable. All ORF alignments show published protein sequences. All ARF alignments show +1 or +2 (*ND7* only) reading frame translations of the full length mRNA sequences. In the ARF alignments of *CR3*, *ND7* and *RPS12*, translations were made using the alternative *T. brucei* mRNA sequences shown in Fig 1. Two alignments of CR3 ARFs are presented to display the *P. serpens* +2 reading frame which has no stop codons. *L. tarentolae* CR4 published protein sequence shows limited homology in the C terminus to all other CR4 protein sequences. The edited mRNA posses two editing sites where 13 U residues are inserted. If the second of these insertion sites is shortened to 12 U residues, the translation of this mRNA has much better homology to other CR4 proteins. Alignments with translations of the two different sequences (13U and 12U) are both shown, with the location of the altered site highlighted in red. While *ND8* does not appear to be dual-coding, this alignment was included as well, for comparison of the conservation of a nondual-coding gene with that of the dual-coding genes. It should be noted that *ND8* is the only nondual-coding gene that is pan-edited in *L. tarentolae*, *L. amazonensis*, and *P. serpens*, and *ND7*, *A6* and *COIII* are only partially edited in these species. The internal M observed in the ND8 sequence is highlighted in yellow (see text). !=Termination codon

**S1 Table.**
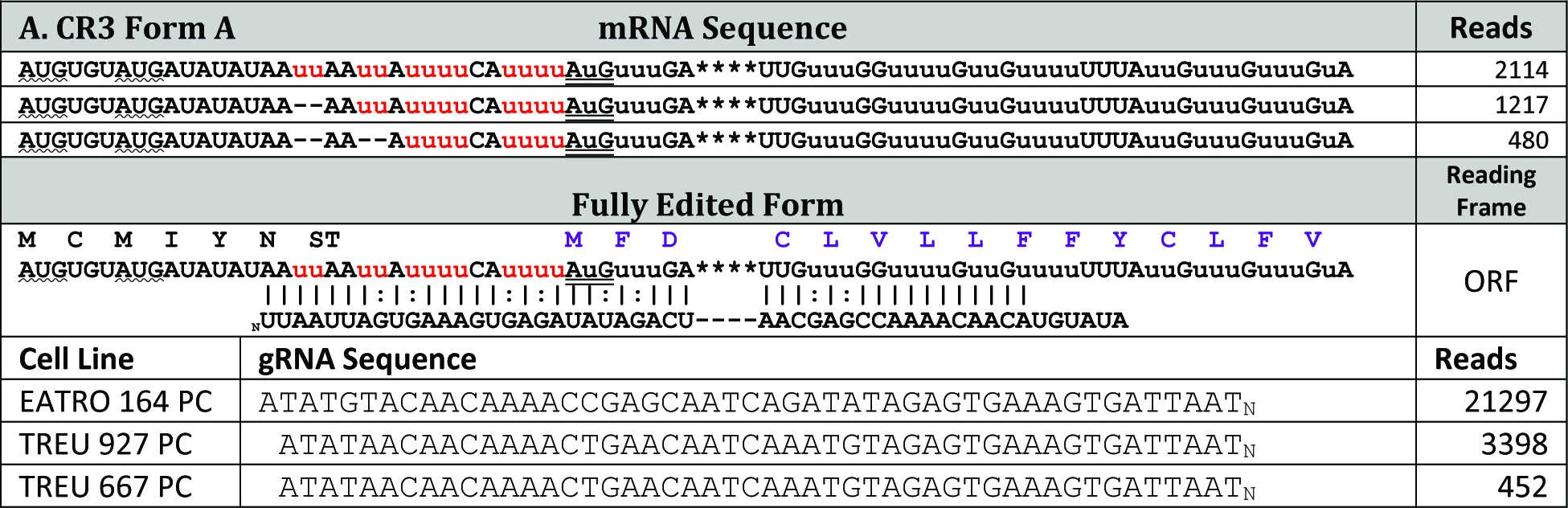
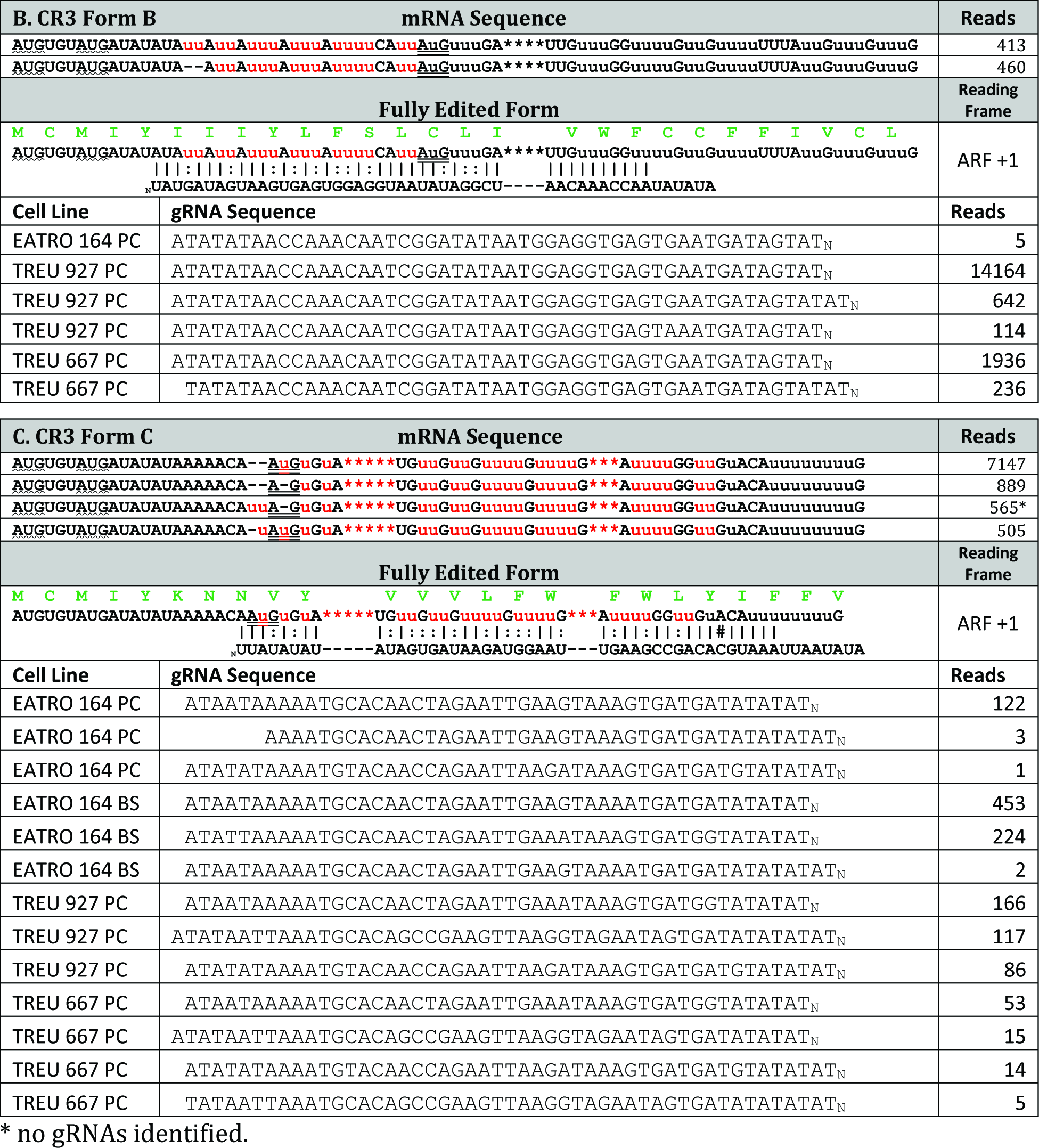
Identified *CR3* mRNA and gRNA transcripts. A - C: Major CR3 mRNA and gRNA sequence classes. The *CR3* mRNA transcriptome was generated using the TREU667 cell line. Identified sequences were then used to search gRNA transcriptomes from four different cell lines: EATRO 164 Bloodstream (BS), EATRO 164 procyclic (PC), TREU 927 procyclic and TREU 667 procyclic. ORF = previously identified Open Reading Frame (purple protein sequence). ARF = Newly identified Alternative Reading Frame (green protein sequence). Alternatively edited nucleotides are shown in Red. Inserted U-residues are lowercase while deleted U-residues are shown as asterisks. Canonical Watson-Crick base pairs (|); G:U base pairs (:). Previously identified start codons are doubled underlined. Potential upstream AUG start codons are indicated by wave underlines. gRNAs were sorted based on guiding sequence class. Sequence variations observed in the 3’-U-tail were ignored in assigning class. Transcript copy number (Reads), were determined by adding all gRNAs of the same sequence class. Only major sequence classes are shown (defined as containing greater than 100 transcript copies). In the case of rare transcripts, the identified gRNA are shown regardless of copy number. gRNA transcript numbers varied greatly between the different cell lines. Interestingly, the most abundant mRNA (*CR3* Form C, 7147 reads) had the fewest identified gRNA reads.

**S2 Table.**
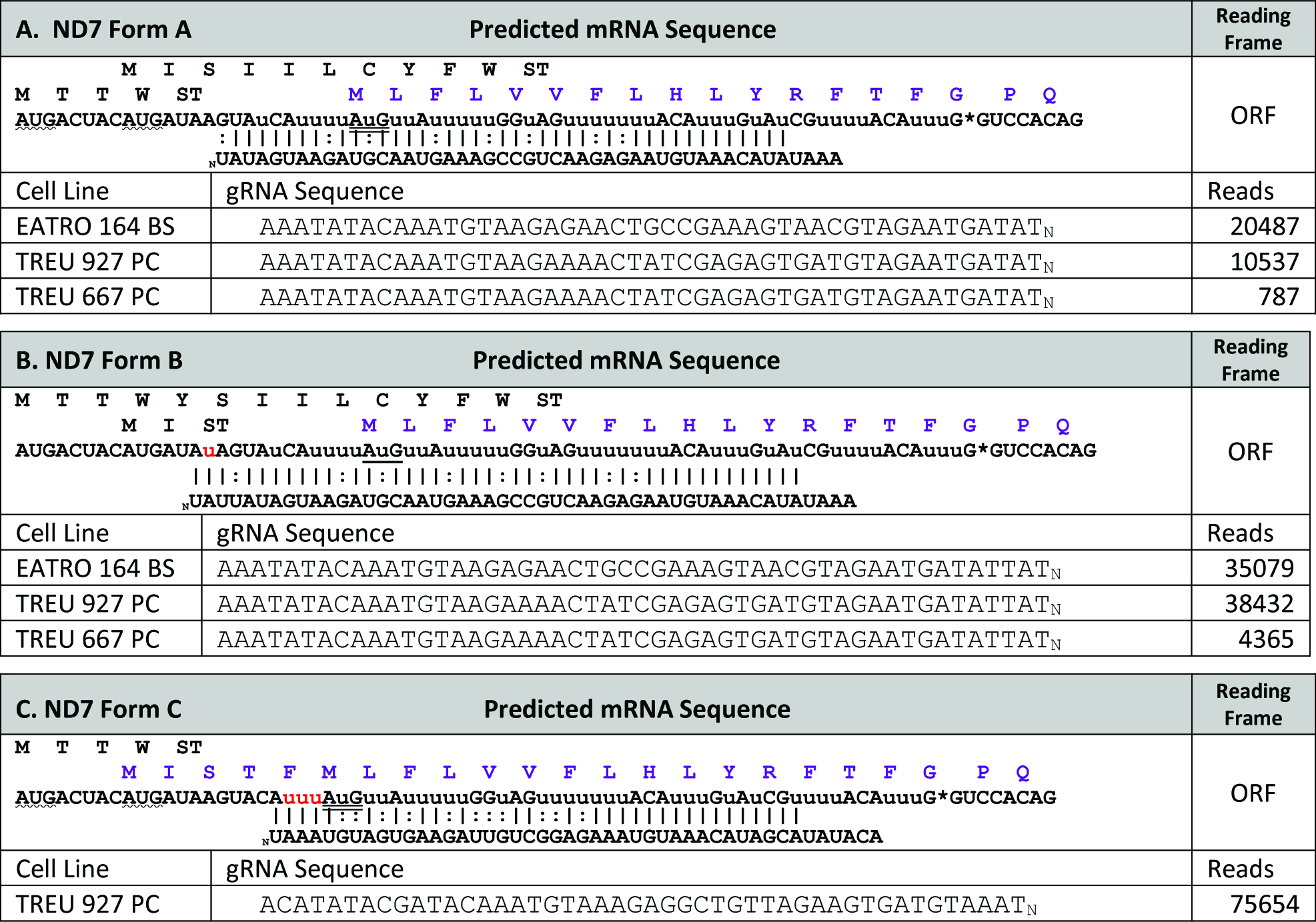
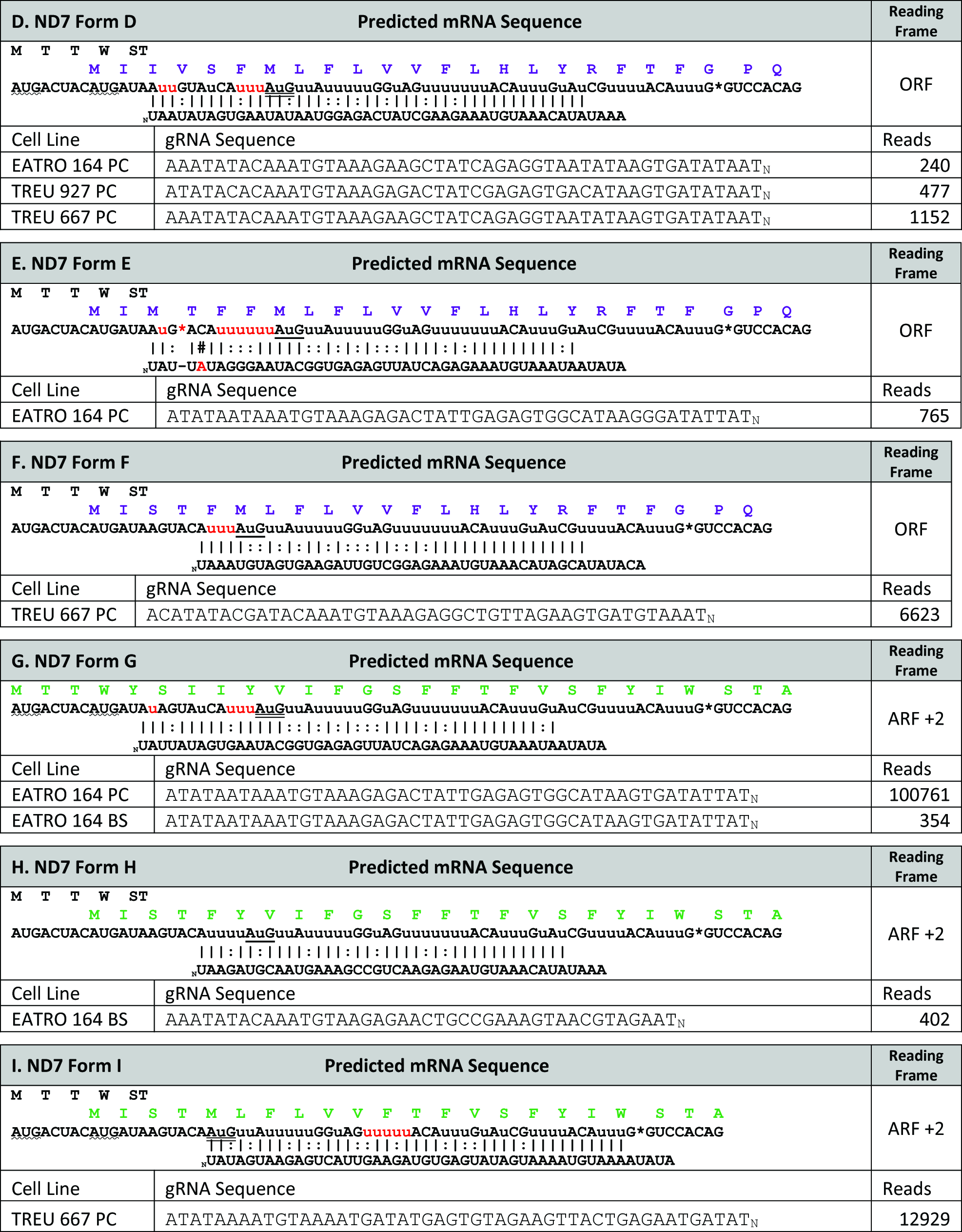
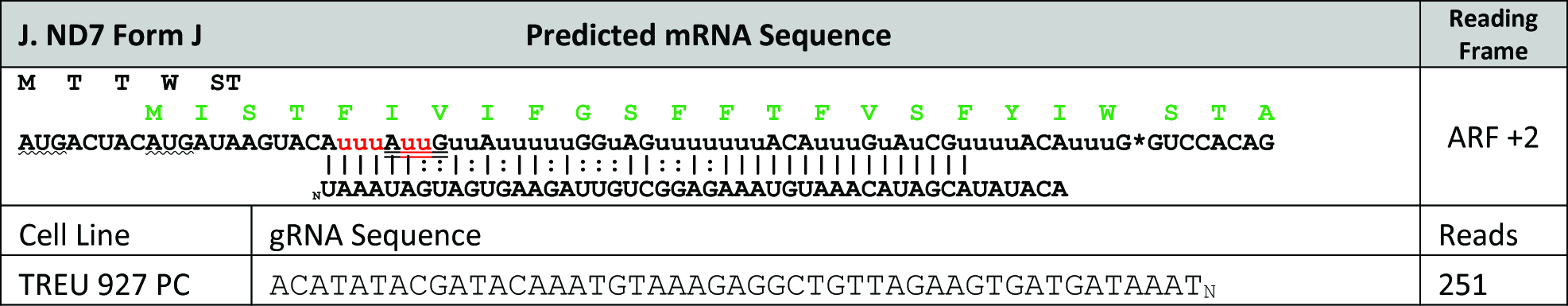
*ND7* 5’-most gRNA populations and the predicted mRNA sequences generated. *ND7* terminal (5’ most) gRNA populations and the predicted mRNA sequence generated. Predicted sequences presented are based on the most abundant gRNAs that generate each reading frame found in the four gRNA transcriptome databases. Initial characterization of the *ND7* transcript was done using the EATRO 164 cell line and is unusual in that it is edited in two distinct domains [20]. While the 5’ domain was edited in both life cycle stages, complete editing of the 3’ domain was only detected in bloodstream stage parasites. Interestingly, the most abundant EATRO 164 PC (procyclic or insect form) gRNA would generate a sequence that brings the 5’ most AUG into a +2 frame. The ARF is 65 AA long and involves the entire 5’ editing domain. In contrast, the most abundant gRNAs in the EATRO 164 Bloodstream stage library (EATRO 164 BS), would generate sequences that use the originally described *ND7* ORF). While gRNA transcript numbers again varied greatly between the different cell lines, all three cells lines had gRNA sequence variants that allowed access to both reading frames.

**S3 Table.**
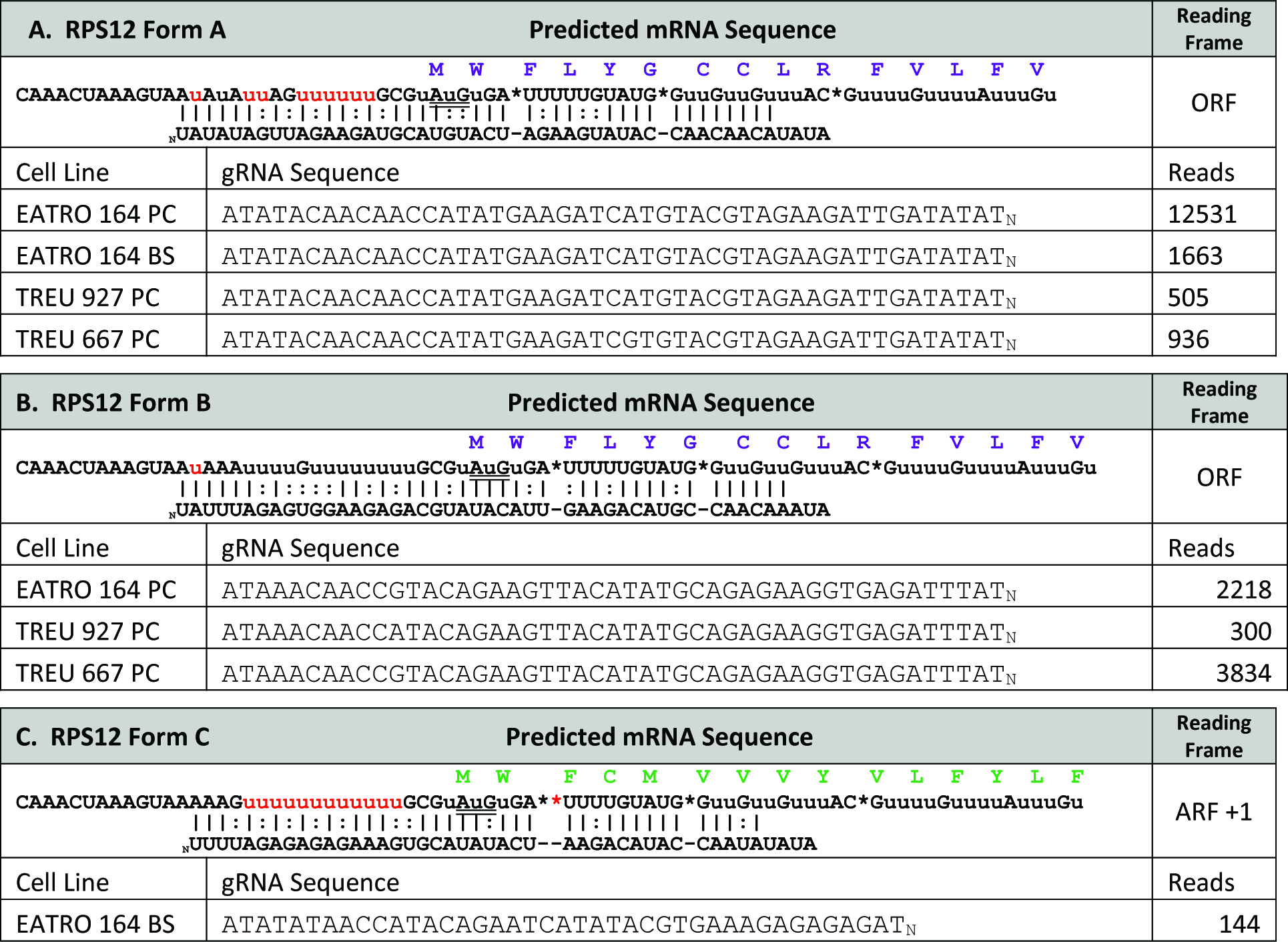
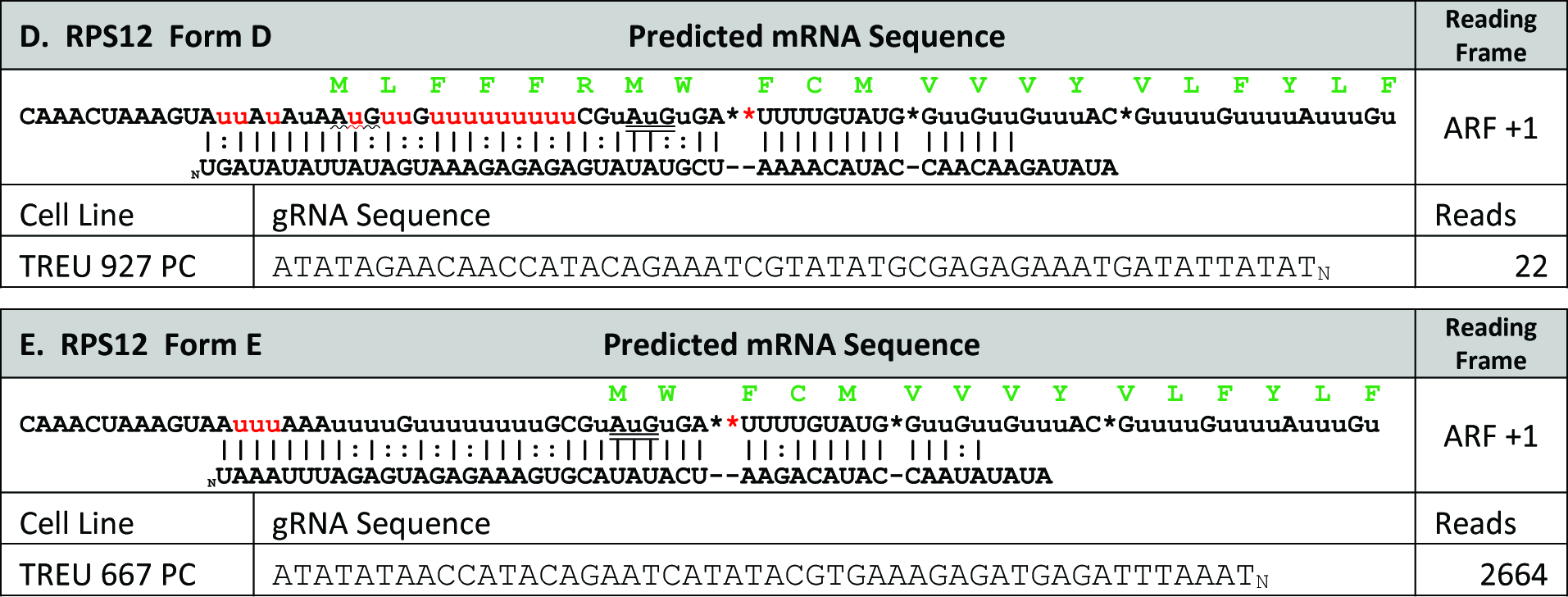
*RPS12* 5’-most gRNA populations and the predicted mRNA sequences generated. *RPS12* terminal (5’ most) gRNA populations and the predicted mRNA sequence generated. *RPS12* differs from both *CR3* and *ND7* in that the alternative edit that shifts the reading frame occurs just downstream of the previously identified start codon (double-underlined). We do note that the identified alternative gRNAs are rare in all of the gRNA libraries except TREU 667.

